# Universal dynamics of cohesin-mediated loop extrusion

**DOI:** 10.1101/2024.08.09.605990

**Authors:** Thomas Sabaté, Benoît Lelandais, Marie-Cécile Robert, Michael Szalay, Jean-Yves Tinevez, Edouard Bertrand, Christophe Zimmer

**Author notes:** These authors contributed equally to the work (Benoît Lelandais, Marie-Cécile Robert). These authors contributed equally to the work (Edouard Bertrand, Christophe Zimmer).

## Abstract

Most animal genomes are partitioned into Topologically Associating Domains (TADs), created by cohesin-mediated loop extrusion and defined by convergently oriented CTCF sites. The dynamics of loop extrusion and its regulation remains poorly characterized *in vivo*. Here, we tracked TAD anchors in living human cells to visualize and quantify cohesin-dependent loop extrusion across multiple endogenous genomic regions. We show that TADs are dynamic structures whose anchors are brought in proximity about once per hour and for 6-19 min (∼16% of the time). TADs are continuously subjected to extrusion by multiple cohesin complexes, extruding loops at ∼0.1 kb/s. Remarkably, despite strong differences of Hi-C patterns between the chromatin regions, their dynamics is consistent with the same density, residence time and speed of cohesin. Our results suggest that TAD dynamics is governed primarily by CTCF site location and affinity, which allows genome-wide predictive models of cohesin-dependent interactions.

## Introduction

Genome-wide chromosome conformation capture methods (such as Hi-C^1^) revealed that mammalian genomes are folded into thousands or tens of thousands of 100-1500 kb large Topologically Associating Domains (TADs)^2–5^. TADs are defined as chromatin regions with a higher frequency of internal contacts as compared to contacts outside their boundaries, and hence appear as blocks on the diagonal of Hi-C contact matrices^2,3^. In Hi-C contact matrices, TADs can display peaks of enhanced contact frequencies at their corners, which are interpreted as signatures of chromatin loops. TADs and loops critically depend on the cohesin complex, since removal of its subunit RAD21 leads to their genome-wide disappearance^6,7^. Both structures are now understood as resulting from loop extrusion, a process wherein the ring-shaped cohesin complex progressively generates a DNA loop of increasing size by consuming ATP^6,8–11^. This process is stopped by convergently oriented CCCTC-binding Factor (CTCF) sites, which act as barriers to cohesin molecules and define TAD boundaries^4,6,12–16^. Loop extrusion is believed to play an important role in key nuclear processes such as gene expression regulation^17–22^, DNA repair^23,24^ and V(D)J recombination^25^. Characterizing the dynamics of cohesin-mediated extrusion is therefore essential to better understand these processes. Critical questions include how frequently any given genomic region undergoes extrusion, how frequently and for how long anchors come into contact, and how rapidly DNA is extruded by the cohesin complex in living cells. It also remains unknown how the dynamics of loop extrusion is regulated across the genome and whether other factors than CTCF or cohesin modulate the dynamics of TADs. Answers to these questions remain fragmentary6,10,11,26,27.

TADs and loops are usually characterized using bulk Hi-C, which yields contact maps that are averaged over millions of cells at a single time point. However, single-cell Hi-C^28^, multiplexed DNA FISH studies^29–31^, single-particle tracking^32,33^ and polymer simulations^9,12^ all revealed that chromatin structure can vary from cell to cell within an isogenic population. Hi-C and DNA FISH are intrinsically limited to fixed cells, and therefore ill-suited to characterize the dynamics of chromatin regions. Experiments that recapitulated loop extrusion *in vitro* have measured cohesin-mediated extrusion rates of 0.5-1 kb/s^10,11^, and thus shed light on some dynamic features of loop extrusion. However, it remains uncertain whether extrusion occurs at a similar, higher or lower rate in living cells, where many molecular factors may slow down or on the contrary accelerate this process. Likewise, numerous factors can potentially tune the loading and release rates of cohesin *in vivo*. For instance, cohesin residence time on chromatin is decreased by WAPL^34–36^ and can be modulated by the STAG subunit of the cohesin complex^37^. A full understanding of loop extrusion therefore requires analyzing this process in single and living cells.

Two recent studies analyzed TAD dynamics in living mouse embryonic stem cells (mESCs) and confirmed that TADs are dynamic structures^26,27^. Being limited to a single locus each, however, these studies did not reveal how extrusion dynamics is regulated across the genome. Additionally, mESCs feature many differences with differentiated cells^38–40^, including less condensed chromatin and a very short G1 phase (∼1-2 hours^41^). Thus, a broader characterization of loop extrusion in single human cells is presently lacking.

Here, we labelled multiple pairs of endogenous TAD anchors in human HCT116 cells, and used live-cell fluorescence microscopy, dedicated analysis methods and polymer simulations to visualize and quantitatively characterize cohesin-dependent loop extrusion. We show that TAD anchors are frequently brought into proximity by loop extrusion but for short time intervals. Moreover, TADs are almost always folded into multiple loops, extruded by several cohesin complexes simultaneously. We also provide the first estimate of loop extrusion speed in living cells. Finally, polymer simulations show that TAD dynamics across different genomic regions are consistent with a narrow range of parameters defining cohesin dynamics (density, residence time and speed), suggesting that extrusion dynamics is universal, rather than locally tuned, and that TAD dynamics is governed primarily by CTCF binding.

## Results

### Visualizing the dynamics of TAD anchors

To visualize and quantify cohesin-mediated chromatin looping in living cells, we labelled endogenous TAD anchors by CRISPR-mediated insertion of TetOx96 and CuOx150 arrays, which are respectively bound by TetR-splitGFPx16-NLS and CymR-NLS-2xHalo (imaged with the bright and photostable dye JFX646^42^; **Figures 1A** and **B**).

**Figure 1:**
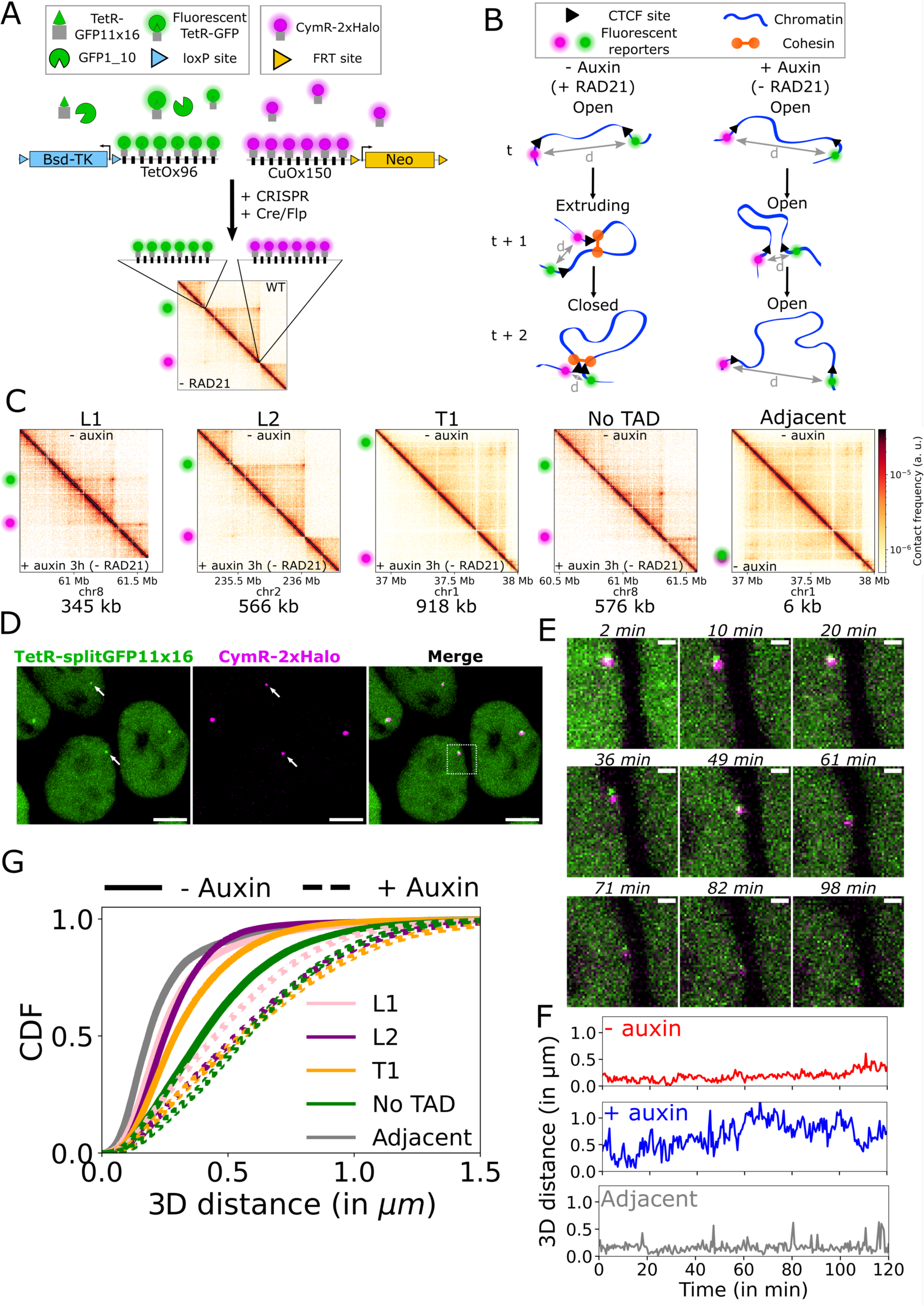
Tracking TAD anchors in living human cells. **A:** TetOx96 and CuOx150 repeat arrays were inserted at TAD anchors in HCT116 cells and visualized using TetR-splitGFPx16 and CymR-2xHalo^JFX646^, respectively. Multimerized GFP11 fragments are not shown for clarity. Antibiotic cassettes (Bsd-TK and Neo) were removed by Cre and Flippase (Flp) recombinases to avoid interference of transcription with loop extrusion. **B:** Cells were imaged with or without auxin treatment, which leads to RAD21 depletion. The 3D distance (d) between the two fluorescent reporters was computed as function of time. In absence of auxin, the chromatin region between the two anchors is in one of three states: open (no loop), extruding (*i.e.* containing one or more DNA loop(s)), or closed (the two anchors are linked by a cohesin complex). **C:** Capture Micro-C maps of cells left untreated or treated with auxin for 3 hours. Green and magenta spots indicate the genomic locations of the inserted repeat arrays. The genomic distance between TAD anchors is indicated below each Micro-C map. The T1 Micro-C map was used to illustrate the Adjacent control. All Micro-C maps show a 1125 kb sized region, at 2 kb resolution. **D:** Live-cell images of L1 TAD anchors at t=0 min. The arrows indicate spots that were identified as not replicated. Scale bar: 5 µm. **E:** Time-lapse images of a magnified region corresponding to the dotted white box in **D**. Scale bar: 1 µm. **F:** Time series of 3D anchor-anchor distances of the L1 locus in untreated cells (red, corresponding to images in **D** and **E**) or auxin-treated cells (blue) and the Adjacent control (grey). **G:** Cumulative distribution function (CDF) of 3D anchor-anchor distances. All images are maximum intensity projections of 20 z-stacks.

Chromatin loops or TADs display a large range of genomic sizes and are characterized by a diversity of patterns in Hi-C maps^43–45^. This diversity results at least in part from variations in the distributions (*e.g.* clustered or evenly spaced), binding and affinities of CTCF sites within TADs, and across the genome^46,47^. To account for this variability, we selected TADs with distinct Hi-C patterns. We chose two TADs of 345 and 566 kb, hereafter called L1 and L2, respectively, and corresponding to TADs with a strong corner peak characteristic of a loop domain. We also selected a 918 kb long TAD, hereafter called T1, without a corner peak (**Figures 1C** and **S1A**). All three domains exhibited SMC1, RAD21 and CTCF ChIP-Seq peaks at both anchors and at least one pair of convergent and bound CTCF sites at their anchors (**Figure S1A**). Within 40 kb around TAD anchors, L1 and L2 displayed only 1 or 2 strong CTCF sites, while the 5’ anchor region of T1 contained 4 weak CTCF sites (**Figure S1A**). Although all three TADs were located within the transcriptionally active A compartment, the L1 and L2 domains exhibited the active H3K27ac histone mark and a weak signal for the repressive H3K27me3 histone mark, whereas the T1 domain was associated with higher H3K27me3 signal (**Figure S1A**). Moreover, these domains did not contain genes or enhancers at their anchors and had little or no gene expression within the domain, thus avoiding potential interferences of transcription with chromatin motion and loop extrusion^48,48,49^ (**Figure S1A**). To further avoid transcriptional interference, we used Cre- and Flippase-mediated recombination to remove the highly transcribed antibiotic resistance genes used to select the integration of repeat arrays (**Figure 1A**). In addition, we generated two control cell lines. In the first one, hereafter called No TAD, the genomic distance between labeled loci was 576 kb, similar to L2, but one of the two labeled loci was located outside of a TAD, far from CTCF sites (**Figures 1C** and **S1A**). Therefore, prolonged cohesin-dependent contacts between these two loci are not expected, making this cell line a negative control for anchors directly linked by a cohesin complex. Conversely, the ‘Adjacent’ control cell line featured two fluorescent reporters genomically adjacent to each other, with a mid-array distance of 6 kb, to serve as an approximate control for anchors in spatial proximity (**Figure 1C**).

We first performed Capture Micro-C and observed that fluorescent tagging of chromatin did not disrupt the formation of TADs (**Figure 1C**). Then, we used spinning disk confocal microscopy to image live cells in 3D, every 30 s during 2 hours (**Figures 1B, D, E**, **S2** and **Video S1**). We removed cells containing replicated spots from the analysis to only examine cells in G1 or early S-phase (**Figures S3E-G**). Using dedicated computational tools, we detected, localized and tracked fluorescent spots, and computed the 3D distance between them as function of time (**Figures 1D-F** and **S3A-D**). Importantly, our localization method computed the fundamental limit to localization precision (Cramér-Rao lower bound^50^) associated to each fluorescent spot at every time point, allowing to take into account localization uncertainties in subsequent analyses. We obtained 150-694 time series per studied locus and experimental condition, totaling n=12,269-93,431 measured distances per condition, with mean localization precisions on distances of 70-105 nm (**Figure S3D** and **Table S1**).

The fluorescent labeling of chromatin loci was performed in a HCT116 cell line containing an homozygote insertion of the auxin-dependent degron mini-AID^51^ to allow rapid depletion of the endogenous RAD21 protein. This system yielded an efficient depletion of 91% after 1 hour of auxin treatment (**Figures S4A-C**) and >94% after 3 hours (**Figures S4D** and **S4E**). We used the AtAFB2 auxin-dependent ubiquitin ligase to limit basal degradation^52,53^ (on average 80% of endogenous RAD21 level remains in untreated cells; **Figures S4D** and **S4F**). Upon auxin treatment, we observed a disappearance of TADs in Capture Micro-C maps, as expected^6^ (**Figure 1C**). Prior to live-cell imaging of TAD anchors, cells were left untreated or pre-treated for 2 hours with auxin. Because cohesin-mediated loop extrusion reduces anchor-anchor distances, auxin treatment was expected to increase these distances. This was indeed the case for all studied loci (**Figures 1G** and **S5A**). Likewise, we observed increased chromatin motion upon RAD21 depletion, in agreement with prior experiments^26,27^ (**Figure S5B**).

Thus, we visualized the dynamics of endogenous chromatin loci in living human cells in presence or absence of cohesin, and precisely computed the 3D distance between TAD anchors as function of time.

### TADs are dynamic structures

At any given time, a pair of anchors is in only one of three possible states: (i) the open state, where the chromatin region between the anchors is free of loops; (ii) the extruding state, where one or more DNA loops are being extruded (actively expanded or temporarily stalled) by cohesin complexes and (iii) the closed state, where the chromatin between anchors is fully extruded and the anchors are maintained in contact by at least one cohesin complex (**Figure 1B**). We first aimed to quantify the fraction, frequency and lifetime of closed states.

Accurately identifying closed states is challenging because stochastic motion of the chromatin polymer can bring the anchors in close proximity even without loop extrusion. Additionally, measuring distances is complicated by noise due to random localization errors and because fluorescent reporters do not directly label CTCF anchors^54,55^. Thus, as a proxy for closed states, we detected temporally sustained intervals in which anchor-anchor distances remained small (**Figures 2A** and **S6**). We hereafter refer to these time intervals as the ‘proximal state’, distinct from the ‘closed state’. To segment time series into proximal states, we used a method previously validated on polymer simulations^54^. This method involves two thresholds: (i) a spatial distance threshold, defined using a theoretical polymer model of the closed state distance distribution (see Methods); and (ii) a temporal threshold, defined as the minimal time interval during which anchor distances remain smaller than the spatial threshold. This temporal threshold was used to minimize false identification of closed states due to stochastic motion of chromatin. Specifically, we set the temporal threshold to 3 min (and also examined variations of its value, see below), which yielded proximal state fractions smaller than 5% in auxin-treated cells, where closed states are not expected (**Figure 2B**). In untreated cells, this procedure led to average proximal state fractions of 17% for L1, 16% for L2, 16% for T1 and 8% for the No TAD control (**Figure 2B**). Upon auxin treatment, the fraction of proximal states decreased by 3.1-3.9 fold, highlighting the importance of cohesin-mediated loop extrusion for the establishment of long-range chromatin interactions (**Figure 2B**). The highest proximal state fraction (44%) was found in the Adjacent control (**Figure 2B**), although distances in this cell line were larger than expected (**Figure S5C**), as already observed in previous locus tracking studies^26,27^.

**Figure 2:**
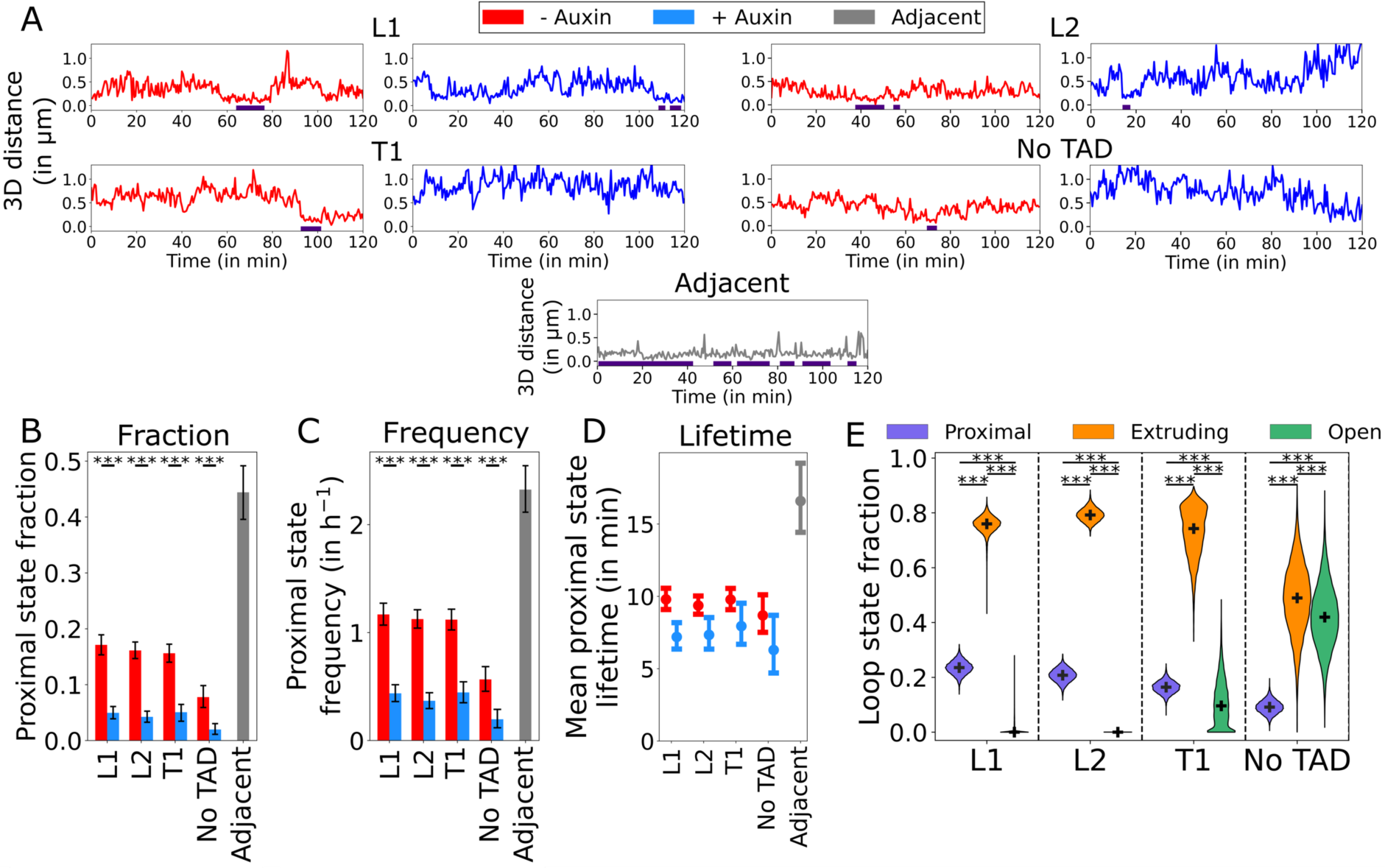
TADs are dynamic structures. **A:** Time series of 3D anchor-anchor distances with or without auxin treatment. The indigo bar indicates segmented proximal states. **B-D:** Fraction (**B**), frequency (**C**) and lifetime (**D**) of proximal states. In **B-C**, data are represented as mean ± 2.5-97.5 percentiles of 10,000 bootstrapped samples ***: P-value < 0.001 from a Mann-Whitney U test. In **D**, data are represented as mean ± 95% CI of the exponential fit used to compute proximal state lifetime. **E:** Fraction of each loop state in untreated cells, estimated from an analytical model of anchor-anchor vectors. The black cross indicates the median and violin plots extend from the minimum to the maximum. ***: P-value < 0.001 from a post hoc Dunn’s test following a Kruskal-Wallis test. N=10,000 bootstrapped samples.

Next, we estimated the frequency of proximal state appearance, *i.e.* the number of transitions to proximal states per unit time. We found that proximal states occurred on average 1.1-1.2 h^-1^ for L1, L2, and T1, 0.6 h^-1^ for the No TAD control and 2.3 h^-1^ for the Adjacent control (**Figure 2C**). Auxin-treated cells exhibited lower frequencies of ∼0.4 h^-1^, consistent with an absence of extrusion (**Figure 2C**). Finally, by exponential fitting of the proximal state durations, we estimated mean proximal state lifetimes of 9.8 min for L1, 9.4 min for L2, 9.8 min for T1 and 8.7 min for the No TAD control (**Figure 2D**), while the Adjacent control exhibited the largest mean lifetimes (16.6 min), as expected (**Figure 2D**). Auxin treatment led to a shortening of proximal state lifetimes by only ∼1.3-fold (**Figure 2D**). This might be a consequence of residual loop formation due to incomplete cohesin degradation (**Figure S4E**).

We assessed how the above estimates depended on the chosen temporal threshold and therefore varied it, obtaining proximal state fractions of ∼1 to 10% in auxin-treated cells. In untreated cells, this resulted in mean proximal state fractions of 8-26%, 8-25%, 7-24%, and 5-12% min for L1, L2, T1 and No TAD, respectively (**Figure S7B**). Therefore, proximal states always represented a minor fraction of loop states regardless of the chosen threshold. The mean proximal state frequency varied in the range 0.3-2.7, 0.3-2.7, 0.3-2.6, 0.2-1.4 h^-1^ for L1, L2, T1 and No TAD, respectively (**Figure S7C**). Thus, proximal states occurred from a few times to a few dozen times per G1 phase. Although mean proximal state lifetimes slightly depended on the chosen threshold (**Figure S7D**), their average values fell consistently within the ranges 6-19 min, 6-16 min, 6-19 min and 4-13 min for L1, L2, T1 and No TAD, respectively (**Figure S7E**).

In summary, we found that in human HCT116 cells, proximal states between TAD anchors occur about 3-27 times during a 10-hour G1 phase, last 6-19 min and yield a total summed duration of about 0.8-2.6 hours in G1.

### TADs are constantly extruded

Having characterized the proximal state, we next quantified the open and extruding states. We used a method previously validated on simulations to quantitatively estimate the fractions of each loop state (closed, extruding and open)^54^. This method uses the total distribution of anchor-anchor vectors and requires knowing the vector distribution in the closed and open states (**Figure S8A**). For the former distribution, we used coordinates belonging to proximal states in untreated cells, whereas the latter distribution was determined from RAD21-depleted cells.

In auxin-treated cells, we estimated 10%, 9%, 11% and 5% of proximal states for L1, L2, T1 and the No TAD control, respectively, similar to the fractions obtained in the previous analysis (**Figures 2B** and **S8B**), and we found 0% of extruding states (**Figure S8B**). Furthermore, we estimated open state fractions of 90%, 91%, 89% and 94% for L1, L2, T1 and No TAD, respectively (**Figure S8B**), consistent with the disappearance of chromatin loops following RAD21 depletion^6^ (**Figure 1C**).

In untreated cells, by contrast, we estimated 24%, 21%, 16% and 9% of proximal states for L1, L2, T1 and No TAD, respectively (**Figure 2E**), in good agreement with the fractions obtained from the analysis above (**Figure 2B**). Most importantly, we estimated extruding state fractions of 76%, 79%, 73% and 49% for L1, L2, T1 and No TAD, respectively. By contrast, the open state was completely absent in L1 and L2 (both medians of 0%) and represented only 11% of loop states in T1, while we found 42% of open state in the No TAD control (**Figure 2E**).

Altogether, these data indicate that, under physiological conditions, TADs are rarely, if ever, in a completely relaxed (open) state but almost permanently undergo cohesin-dependent loop extrusion.

### TAD anchors are brought together at rates of 0.06-0.2 kb/s in living cells

Next, we aimed to determine the speed at which TAD anchors are brought together. We hereafter distinguish two quantities, both expressed in bp/s: (i) the molecular speed at which the cohesin motor pulls out strands of DNA to form a loop (hereafter referred to as ‘motor speed’), and (ii) the rate at which the unextruded DNA between the anchors is reduced, hereafter called ‘closing rate’ (**Figure 3A**). For a single cohesin complex extruding DNA between two anchors devoid of obstacles, the closing rate equals the motor speed (**Figure 3A**). However, in the presence of multiple cohesin complexes and internal CTCF sites where cohesin can stall, the closing rate can be either smaller or larger than the motor speed. Neither quantity has been measured in living cells to date, while motor speeds of 0.5-1 kb/s were estimated *in vitro*^10,11^.

**Figure 3:**
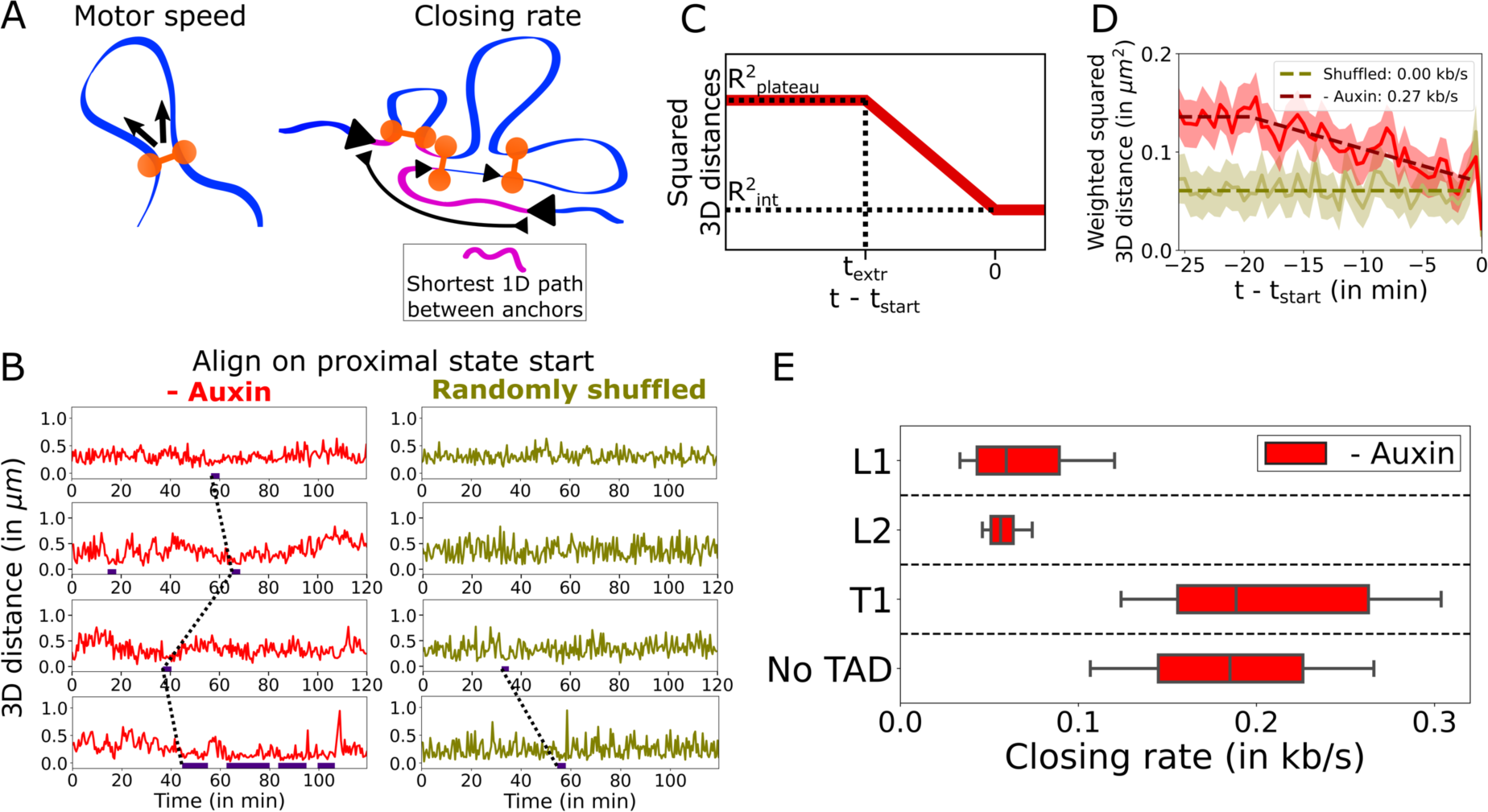
TAD anchors are brought together at rates of ∼0.1 kb/s in living cells. **A:** Cohesin motor speed and closing rate are two different measures of extrusion speed. The cohesin motor speed is the number of DNA base pairs extruded by a single cohesin complex per unit time on DNA devoid of obstacles (*e.g.* CTCF sites). The closing rate measures the rate at which the effective loop-free DNA between TAD anchors, *i.e.* the shortest 1D path between anchors (pink), decreases as a result of extrusion. The closing rate reflects the action of single or multiple cohesin complexes extruding simultaneously, as well as cohesin stalling at internal CTCF sites. Cohesin molecules not involved in the shortest 1D path do not contribute to the closing rate. **B:** Distance time series of the T1 locus without auxin treatment before (left) and after (right) random shuffling of timepoints. Indigo bars indicate segmented proximal state intervals. Dotted lines indicate alignment of the starting times *t*_start_ of proximal states across single time series. **C:** Fitting strategy to determine closing rates. A piecewise linear model with three parameters 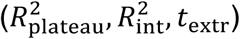 is fitted to the mean squared distances as function of time after alignment to *t*_*start*_. The slope of the linear decrease defines the closing rate. **D:** Aligned time series of mean squared 3D distances measured on the T1 locus, weighted by localization precisions, before (red, N=114) or after (dark yellow, N=79) random shuffling. The linear fit is shown as a dashed line. The shaded area represents the weighted SEM. **E:** Estimated closing rate on time series from untreated cells. Boxplot whiskers extend from 10 to 90 percentiles of n=5,000 bootstrapped samples.

We first focused on measuring the closing rate. This task is challenging because the progressive decrease in anchor-anchor distance expected from extrusion can be obscured by stochastic chromatin motion compounded by random localization errors^54,55^. To reduce the effect of these fluctuations, we averaged together many single time series, as previously validated on polymer simulations^54^. We reasoned that timepoints immediately preceding the closed states should be undergoing extrusion, and therefore aligned distance time series from a large number of cells on the starting time of proximal states^54^ (*t*_*start*_; **Figures 3B-D**). To ensure that we considered exclusively timepoints in the extruding state, we ignored proximal states preceded by another proximal state (**Figure 3B**). In order to estimate the closing rate, we then fitted to the aligned and averaged squared distances a 3-parameter model consisting of a constant plateau followed by a linear decrease, using a weighing scheme to account for variable localization precision (**Figure 3C**, see Methods). The slope of the linear decrease directly provided an estimate of the closing rate. In order to determine if the data actually reveal this linear decrease, we also fitted a constant (1-parameter) model and selected the model with the lowest Bayesian Information Criterion^56^ (BIC; **Figure S9A**, see Methods).

By definition, anchor-anchor distances preceding proximal states are larger than during proximal states, potentially leading to a bias in closing rate estimation. Therefore, as a control, we randomly shuffled all time points within single time series, thereby destroying any signature of processive dynamics (**Figures 3B** and **3D**). Randomly shuffled time series were mostly flat, consistent with a constant model of distances in ∼35% of bootstraps and associated with median closing rates of 0 kb/s (**Figures 3D** and **S9A-C**). By contrast, all untreated time series showed the expected linear decrease of squared distances before *t*_*start*_, as reflected by the selection of the piecewise linear model in 100% of bootstraps (**Figures 3D**, **S9A** and **S9B**). Therefore, our analysis captured the expected linear decrease in squared distances expected from the processive dynamics of loop extrusion in untreated cells, while this behavior was not identified in randomly shuffled time series, as expected. In untreated cells, this analysis yielded closing rates of 0.06 kb/s for both L1 and L2 and of 0.19 kb/s for T1 and No TAD (**Figure 3E**).

As mentioned above, the relation between closing rate and cohesin motor speed is complex, because the former also depends on the locus-specific location and binding affinity of CTCF sites. We therefore turned to polymer simulations to estimate the motor speed of cohesin, and to better understand the impact of cohesin dynamics parameters on TAD structure and dynamics.

### Polymer simulations of cohesin- and CTCF-dependent TAD dynamics

Polymer simulations can predict contact maps and chromatin dynamics based on a small number of parameters^43^. We asked how cohesin dynamics, together with the known locations and affinities of CTCF sites, affect contact patterns and the dynamics of TAD anchors.

For this purpose, we simulated a 2.6 Mb region centered on the TAD anchors of each studied genomic locus, accounting for the dynamic binding and unbinding of cohesin and CTCF to chromatin. Cohesin dynamics was defined by cohesin density, residence time and motor speed (**Figure 4A**). Our simulations allowed multiple cohesin to extrude loops simultaneously, and assumed a 50% probability of cohesin stalling upon CTCF site encounter, consistent with the 50% occupancy of CTCF site estimated in mESCs^57^. CTCF ChIP-Seq peaks were used to determine the local residence time of CTCF at each binding site (**Figure 4B**, see Methods), assuming a median genome-wide residence time of 2.5 min, based on estimates of 1-4 min in mESCs^32^. We independently varied cohesin residence time from 2 to 33 min, cohesin density from 1 to 40 per Mb and simulated cohesin motor speeds of 0.25 or 1 kb/s. These ranges encompassed previous experimental or computational estimates^9,26,27,32,36,57–61^.

**Figure 4:**
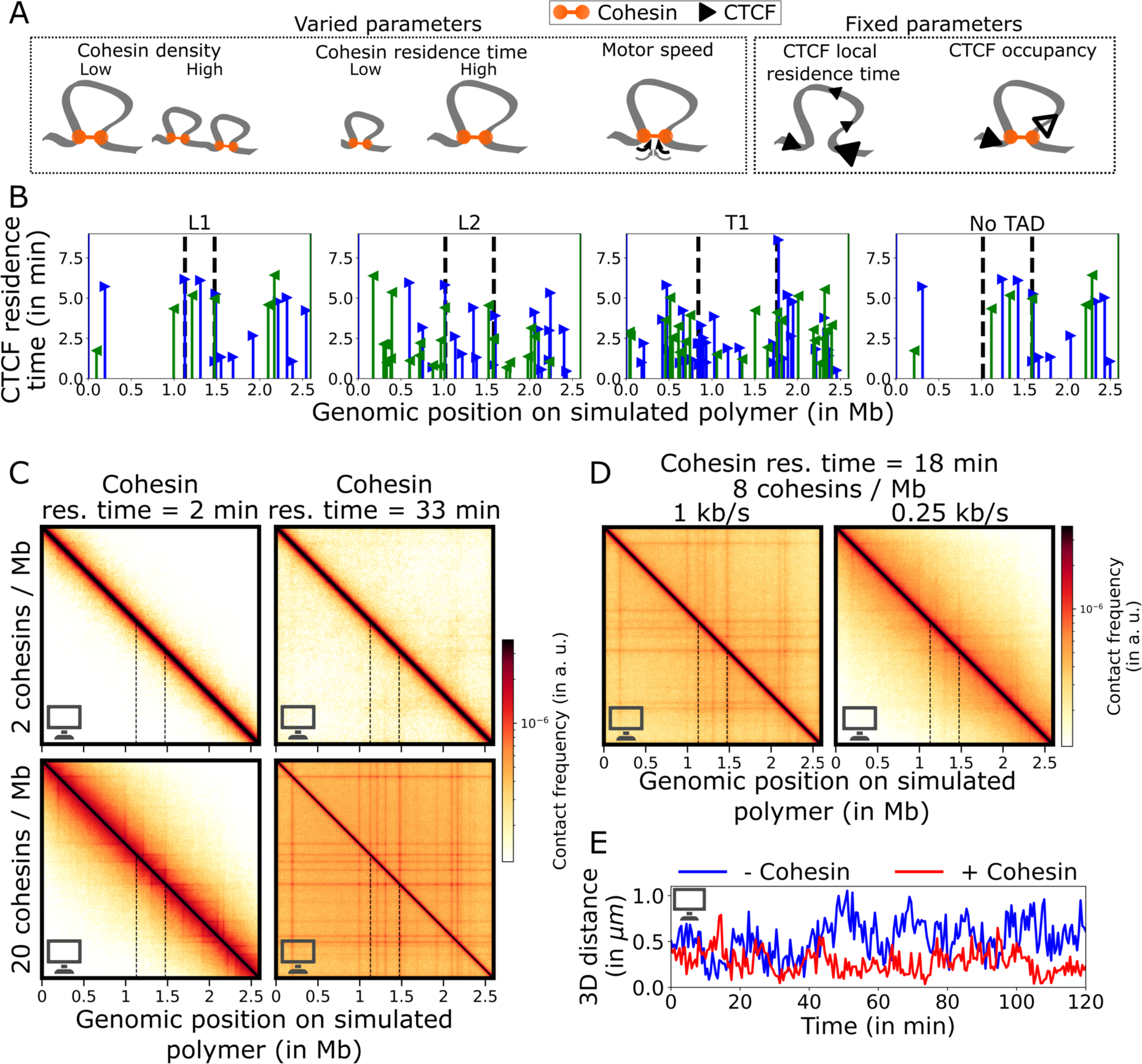
Polymer simulations of cohesin- and CTCF-dependent TAD dynamics. **A:** Parameters used to model loop extrusion in polymer simulations. Three parameters characterizing cohesin dynamics (density, residence time and motor speed) were varied systematically, while the locations, affinities and occupancy of CTCF sites were held constant. **B:** CTCF residence times used for simulations at the four genomic regions. The color and orientation of arrowheads indicates CTCF site orientation. **C-D:** Contact maps from polymer simulations of the L1 locus for different combinations of cohesin residence time and density, for a speed of 1 kb/s (**C**) and for speeds of 0.25 kb/s and 1 kb/s (**D**). In **B-D**, black dotted lines indicate the location of TAD anchors (or of fluorescent reporters for the No TAD control). **E:** Simulated 3D distance time series for the L1 locus. Simulations include random localization errors (increasing over time and consistent with experimental photobleaching) and reporter-anchor separations.

From the ensemble of simulated polymer conformations, we computed contact maps and averages of contact frequencies as function of genomic distance, p(s). Depending on the assumed parameters of cohesin dynamics, these simulated contact maps featured TADs, corner peaks and stripes (**Figure 4C**). As expected, longer cohesin residence times led to longer loops and flatter p(s) curves, while higher cohesin densities yielded multiple sharp stripes and increased overall contact frequencies (**Figures 4C** and **S10A**). Decreasing cohesin motor speed from 1 to 0.25 kb/s, yielded shorter loops because for a constant residence time, the genomic processivity was shorter (**Figure 4D**). Also, cohesin complexes spent on average less time stalled at CTCF sites relatively to actively extruding, resulting in weaker contacts at CTCF sites (**Figure 4D**).

Next, we used the same simulations to generate time series of anchor-anchor distances (**Figure 4E**). To facilitate comparison with experimental data, we added known reporter-anchor separations and random localization errors (consistent with experiments and increasing with time as a consequence of photobleaching, see Methods). Even in absence of extrusion, anchors occasionally came into close proximity for short periods of time due to stochastic polymer motion (**Figure 4E**). Increasing the density of cohesin led to smaller average distances between anchors, and 6 cohesin complexes per Mb were sufficient to reduce anchor distances by ∼50% on average (**Figure S10B**). The collective action of multiple cohesin complexes therefore greatly contributed to extrusion-dependent shortening of distances between TAD anchors.

Increasing cohesin density led to multiple loops simultaneously constraining anchor motion, as reflected by a decrease in the plateau of 2-point MSD curves (**Figure S10C**), and in agreement with previous studies^26,27^. Interestingly, cohesin residence time had a non-monotonous effect: lengthening residence time from 2 to 12 min (implying cohesin processivities from 90 to 500 kb at 1 kb/s) constrained anchor motion, but lengthening it further (to 33 min, implying a processivity of 978 kb at 1 kb/s) increased anchor motion (**Figures S10D** and **S10E**). At intermediate residence times, cohesin complexes reached TAD anchors and constrained their motion, whereas shorter residence times led cohesin complexes to fall off before reaching TAD anchors, leaving their motions unaffected. For longer residence times cohesin complexes continue past the anchors, again leaving them more often unconstrained (**Figure S10E**).

Note that closed states were rare even at the highest cohesin density of 40 Mb^-1^, for which closed state fractions did not exceed 12%, 4% and 2% for L1, L2 and T1, respectively (**Figure S10F**).

Polymer simulations thus allow to predict the complex dependence of TAD dynamics on cohesin residence time, motor speed and density, together with known CTCF site locations and affinities. We next proceeded to compare these predictions to experimental imaging and Hi-C data in order to determine the cohesin dynamics parameters consistent with experiments.

### TAD dynamics is consistent with universal cohesin dynamics and is predicted by CTCF sites only

Our data on multiple genomic regions allowed us to determine whether the parameters of cohesin dynamics vary across the genome. Using polymer simulations, we determined the cohesin parameters consistent with both live-cell imaging and Micro-C data, first considering the four chromatin regions independently. Any differences in the estimated cohesin parameters across genomic domains may indicate locus-specific and CTCF-independent regulation of loop extrusion dynamics, while an absence of differences would suggest universal cohesin dynamics across the genomic regions.

For Micro-C data, we focused on the p(s) curve and for imaging data, we considered the 2-point MSD curves, the anchor-anchor distance distributions, as well as the fractions of segmented proximal states, their frequencies and lifetimes (**Figure S11A**). We first varied the cohesin density and residence time, while fixing the cohesin motor speed at 1 kb/s. We computed the deviation of polymer simulations from experimental data independently for each of the four loci and highlighted the 10% and 25% best sets of parameters (*i.e.* parameters that minimized this deviation; **Figures 5A** and **S11A**, see Methods). Remarkably, for all four loci, only a relatively narrow region of parameters was consistent with experiments. For residence times exceeding 8 minutes, the 10% best parameter sets comprised cohesin densities between 8 and 14 Mb^-1^ for all chromatin regions (**Figure 5A**). Shorter residence times required much higher cohesin densities (>14 Mb^-1^) to recapitulate experimental observations (**Figure 5A**).

**Figure 5:**
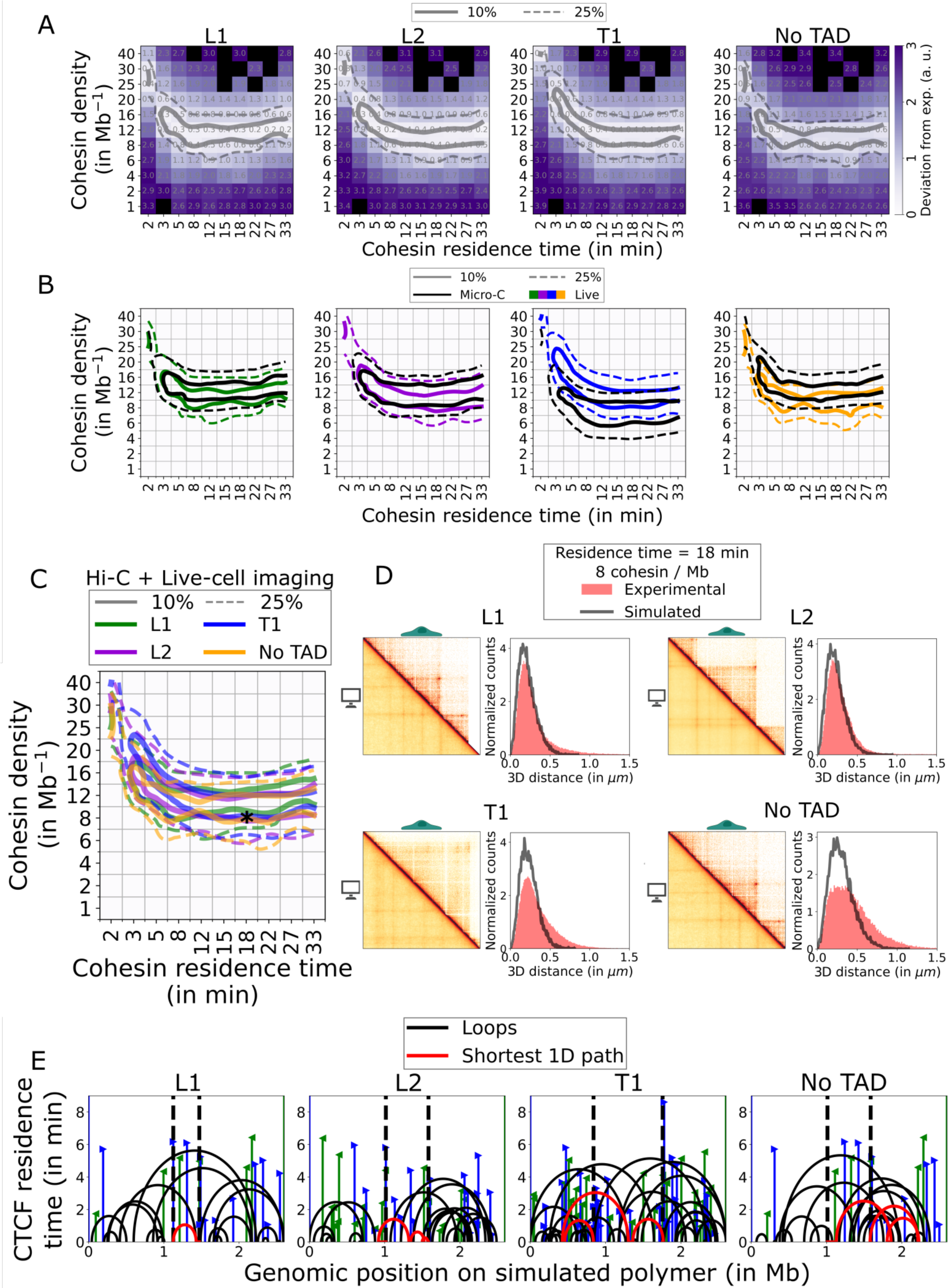
TAD dynamics is consistent with universal cohesin dynamics and is governed by CTCF sites only. **A:** Deviation of polymer simulations from experiments, assuming a cohesin motor speed of 1 kb/s. Black squares correspond to non-assessed parameter combinations. **B:** Contour plots of the deviation of simulations from experiments considering separately Micro-C data (black) or live-cell imaging data (colors). **C:** Superposed contour plots for all genomic regions, based on Micro-C and live-cell imaging data taken together. In **A-C**, solid and dashed lines indicate the 10% and 25% best parameter sets, respectively. **D:** Contact maps and distance distributions of experimental data and simulations. Contact maps display a 1125 kb region centered around TAD anchors, at 2 kb resolution. **E:** Snapshots of 1D representations of extruding loops. The red continuous line shows the shortest 1D path between TAD anchors. The black vertical dotted lines indicate the location of TAD anchors. Simulations used for **D** and **E** assumed the combination of parameters highlighted by a black asterisk in **C**.

These results were consistent even when separately considering experimental Micro-C and live-cell imaging data. For TADs L1 and L2, the 10% and 25% best parameter combinations inferred from Micro-C data were in excellent agreement with those inferred from imaging data (**Figure 5B**). For the T1 and No TAD regions, Micro-C and imaging data were in good agreement since the 10% best sets overlapped for a narrow range of parameters (9 Mb^-1^ and 11 Mb^-1^, respectively) and the 25% best sets of parameters overlapped for at least half of the parameter sets (**Figure 5B**). Thus, our comparisons indicate a high consistency between parameters estimated from Micro-C data of fixed cells and imaging data of living cells separately, thereby providing reciprocal validation of these estimates from two very different experimental techniques.

We then compared the cohesin parameters inferred independently from the four regions, considering both Micro-C and imaging data together. Strikingly, the inferred parameter ranges were largely in agreement across all regions, as observed by the overlap of the parameter space consistent with experiments (**Figure 5C**). For the four chromatin regions, the 10% best sets of parameters overlapped for cohesin densities of 7-12 Mb^-1^ and residence times longer than 8 min, or cohesin densities of 12-16 Mb^-^1 and residence times of 3-8 min (**Figure 5C**). Applying the same analysis to auxin-treated cells led to a decrease of 2 fold in the estimated cohesin density (**Figure S11B**).

To gain further insights into chromatin dynamics consistent with our data, we fixed the cohesin density to 8 Mb^-1^ and the residence time to 18 min, a combination of parameters that allowed a good match of simulations with experimental data for all four regions (**Figure 5C** and **Video S2**). Using these parameters, we could reproduce experimental contact maps (**Figure 5D**) and P(s) curves (**Figure S11C**), 2-point MSD curves (**Figure S11D**) and distance distributions (**Figures 5D** and **S11E**) for L1, L2 and T1 at once, although the agreement with experimental data was poorer for the No TAD region. This combination of parameters led to closed state fractions of 2.7%, 1.1% and 0.6% for L1, L2 and T1, respectively (**Figure S10B**), indicating that direct cohesin-dependent anchor-anchor contacts are rare.

In order to estimate the number of loops connecting the anchors, we computed the shortest 1D path of DNA between TAD anchors (**Figures 3A** and **5E**). On average and at any timepoint, TAD anchors were connected by 2.4-2.8 internal loops (**Figures 5E** and **S12A**, **Video S2**) and separated by effective DNA path lengths of 144, 142 and 135 kb for L1, L2 and T1 (**Figure S10H**). Thus, although the TADs greatly differed in size (by 2.7 fold from 345 to 918 kb), these DNA path lengths varied by only 7%, which suggests that loop extrusion homogenizes effective genomic separations. Counter-intuitively, we noticed that DNA sequences around, and not only within, the TAD anchors could also be involved in the shortest 1D path (**Figure 5E**). Thus, the dynamics of TADs can be influenced by CTCF sites and cohesin complexes located at large genomic distances (up to 800 kb) from TAD anchors. This highlights the need to take into account neighboring regions when assessing cohesin-dependent chromatin interactions of a specific locus.

### Cohesin extrudes loops at ∼0.1 kb/s in living cells

Decreasing the cohesin motor speed from 1 to 0.25 kb/s in simulations did not yield appreciable differences in deviations from experiments (minimal deviations of ∼0.3 for both motor speeds; **Figures S11F** and **S11G**). To estimate the cohesin motor speed, we therefore turned to a different approach.

We computed closing rates from polymer simulations with varying cohesin motor speeds, using the same procedure as for experimental data (**Figures 3**, **6A** and **6B**). To do so, we fixed the cohesin density and residence time to 8 Mb^-1^ and 18 min, respectively, and simulated motor speeds of 0.125, 0.25, 0.5 and 1 kb/s. Using the same fitting method as above, we could detect the expected decrease of distances due to extrusion in all the simulated conditions, as compared to a constant model (**Figure S13A**). Noteworthy, because different TADs exhibit different distributions of CTCF sites and different genomic lengths, the same motor speed could yield different closing rates on distinct TADs (**Figure 6C**). Additionally, alignment of time series on proximal states tended to underestimate closing rates, as compared to time series aligned on closed states (**Figures S13B** and **S13C**). Critically, however, cohesin motor speeds below 0.25 kb/s could simultaneously explain the different closing rates derived from experimental data for L1, T1 and No TAD, whereas the other tested speeds could not (**Figure 6C**). Specifically, L1 and No TAD regions were consistent with cohesin motor speeds below 0.125 kb/s, while T1 was consistent with motor speeds slightly above 0.125 kb/s and below 0.25 kb/s. The L2 region behaved differently and was consistent with motor speeds exceeding 1 kb/s. Unexpectedly, increasing cohesin motor speed decreased the estimated closing rates for L2, T1 and No TAD (**Figure 6C**). This emerged from the larger distances observed at lower cohesin motor speeds, which increased the estimated closing rates (**Figure 6A**). Thus, our analysis indicates that cohesin motor extrudes DNA at rates of ∼0.1 kb/s in living cells, 5-10 fold slower than *in vitro*^10,11^. This constitutes the first measurement of cohesin-dependent extrusion speed in living cells.

**Figure 6:**
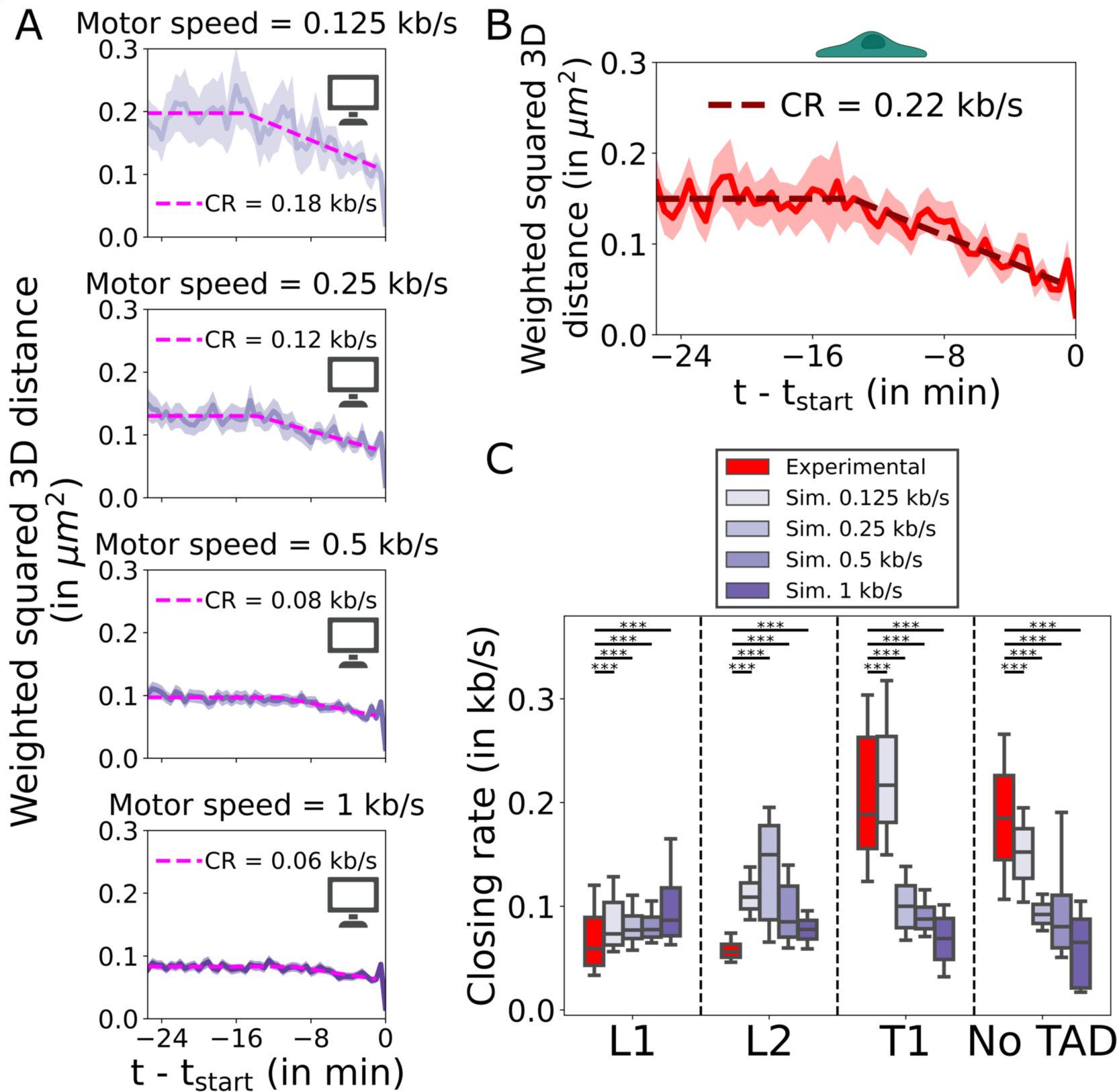
Cohesin extrudes loops at ∼0.1 kb/s in living cells. A-B: Average squared distance time series, weighted by localization precision, aligned on *t*_*start*_ and the corresponding piece-wise linear fit for closing rate (CR) estimation (dotted) for simulated (**A**) and experimental (**B**) data of the T1 locus. In **A**, the simulated cohesin motor speed is indicated above each condition. In **A** and **B**, the shaded area indicates the weighted SEM. **C:** Closing rates determined from experimental (red) or simulated (shades of purple) distance time series. Simulations assumed a cohesin motor speed of 0.125, 0.25, 0.5 or 1 kb/s. Boxplot whiskers extend from 10 to 90 percentiles of N=5,000 bootstrapped samples, N=600 distance time series were used for each simulated condition. ***: P-value<0.001 from a Mann-Whitney U test, adjusted for multiple testing by Bonferroni correction. Simulations assumed a cohesin density of 8 Mb^-1^ and a residence time of 18 min.

Together, our simulations of multiple endogenous loci argue against strong variations in cohesin density and instead suggest that this parameter is universal across the genome. While we cannot rule out large differences in residence times across loci, our data are also consistent with a unique cohesin residence time across the genome. Thus, the observed variations in TAD dynamics and contact maps between the four chromatin regions can be explained without major changes in the three parameters of cohesin dynamics (density, residence time and motor speed), with the exception of the motor speed in the L2 TAD. Instead, they can be explained as a result of the different locations and binding strengths of CTCF sites only. In summary, our data suggest that cohesin dynamics is universal rather than locally regulated and that the variations in TAD dynamics are determined by CTCF binding.

## Discussion

We used live-cell microscopy to track the motion of endogenous TAD anchors at multiple genomic regions in human HCT116 cells, in presence or absence of cohesin. We quantitatively characterized cohesin-dependent DNA loop extrusion and used polymer simulations to determine the parameters of cohesin dynamics that govern this process.

First, our data show that TADs are highly dynamic structures and that anchor-anchor contacts are transient, thereby extending to human cells claims initially made on mESCs^26,27^. Therefore, cohesin-mediated long-range interactions are also short-lived in human cells, despite having G1 phases ∼10-fold longer than mESCs. Our analysis indicates that TAD anchors are in spatial proximity for 7% to 26% of the time (**Figure 2B**). Moreover, our simulation-based predictions of closed state fractions ranged from 0.6 to 2.7% for TADs of 345-918 kb (**Figure S10F**). This shows that TAD anchors are in spatial proximity for a significant amount of time but that direct anchor-anchor contacts are very rare. Previous estimates of the fractions of time in the closed state in mESCs ranged from 2-3% for an endogenous 505 kb TAD^26^ to 20-31% for a strong synthetic 150 kb TAD^27^. We further estimated that proximal states have average durations of 6-19 minutes (**Figures 2D** and **7**). Despite the differences in species and analysis methods, our estimates of the duration of proximal states are in remarkable agreement with mean lifetimes of 5-45 min estimated in mESCs^26,27^. While proximal states are transient, our data indicate that they are relatively frequent, since we estimated that TAD anchors come into proximity 0.3 to 2.7 times per hour, *i.e.* 3 to 27 times during a single 10-hour long G1 phase, on average (**Figures 2C** and **7**). Thus, our results establish the highly dynamic nature of TADs in human cells, which suggests that the dynamic process of loop extrusion itself is more functionally relevant than anchor-anchor interactions.

**Figure 7:**
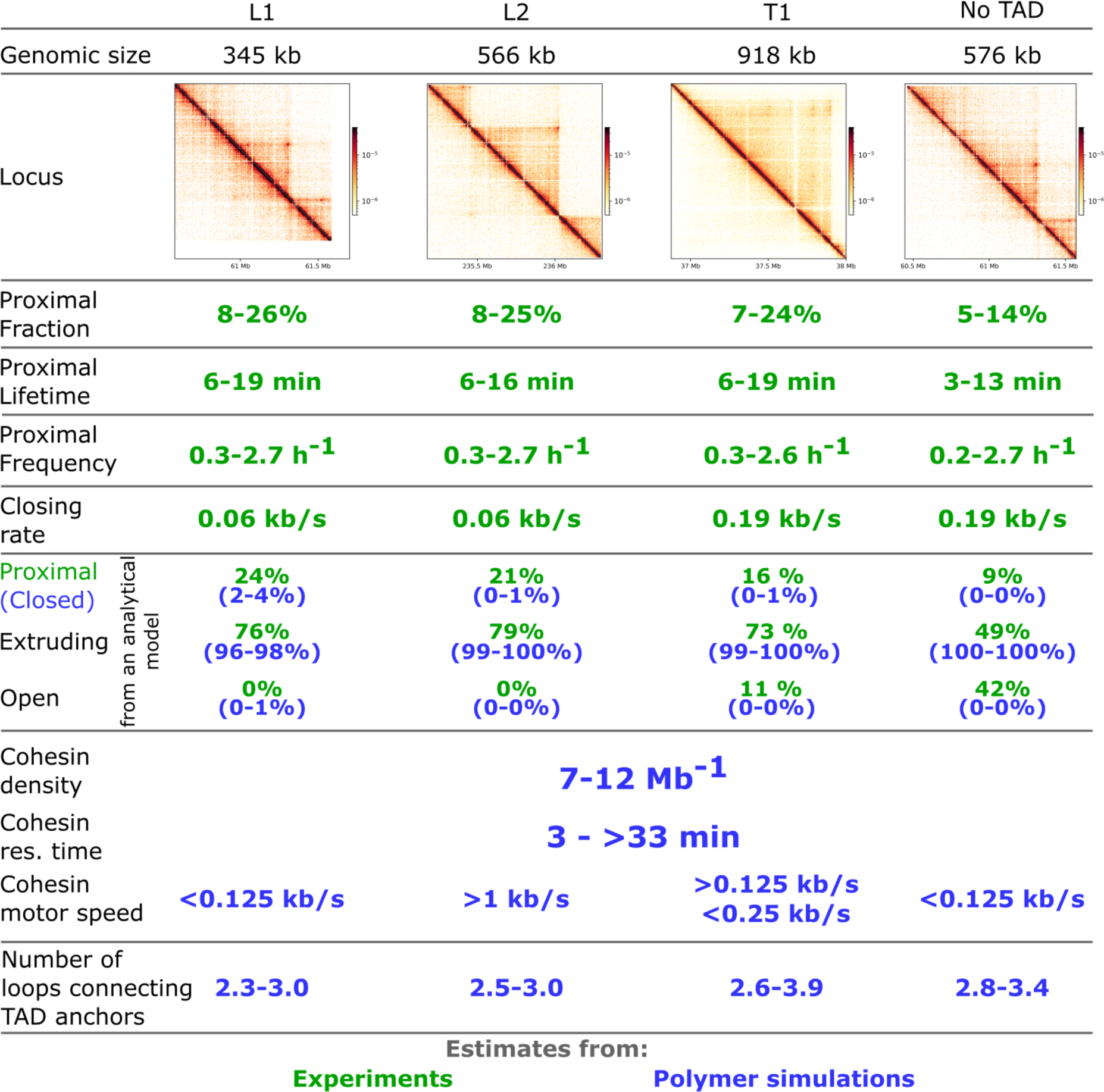
Summary of findings on TAD dynamics at multiple regions of the human genome. Values obtained from a direct analysis of experimental data are shown in green, while values estimated using polymer simulations are shown in blue, with ranges indicating the minimum and maximum values for the 10% best parameter sets across all loci.

Second, we found that the examined TADs (of 345, 566 and 918 kb) are most often in a partially extruded state and almost constantly undergoing loop extrusion but rarely, if ever, in a purely open state (0%, 0% and 11% for L1, L2 and T1, respectively; **Figure 2E**). This is consistent with a previous study in mESCs^26^, and argues in favor of a model where TADs emerge from a collection of growing loops^62^, rather than from a single loop. Therefore, it is likely that several cohesin complexes are simultaneously bound within a single TAD. Assuming that the smallest TAD (L1, with a size of 345 kb) contains at least two cohesin complexes yields a minimum cohesin density of 5.8 Mb^-^1, in agreement with the lower bound of 7 Mb^-1^ estimated from our polymer simulations (**Figure 5C**).

Third, our study provides the first estimate of the speed of DNA loop extrusion *in vivo*. We distinguished the cohesin motor speed from the closing rate, *i.e.* the rate at which the effective genomic separation between TAD anchors diminishes. We estimated closing rates of 0.06-0.19 kb/s from live-cell imaging (**Figure 3E**), and a cohesin motor speed of ∼0.1 kb/s for three out of the four examined genomic regions, from comparison of polymer simulations to experiments (**Figure 6C**). This estimated speed of 0.1 kb/s is 5-10 fold lower than estimates of 0.5-1 kb/s from *in vitro* experiments with purified proteins^10,11^. Consequently, cohesin-mediated loop is slowed down in cells, possibly by the crowded chromatin environment (*e.g.* nucleosomes or replication machineries acting as roadblocks^63,64^) or by other nuclear factors directly reducing its motor speed. This is close to the estimate of ∼0.2 kb/s obtained when dividing 230 kb (the genome-wide median loop length, **Figure S1B**) by the average cohesin residence time of 20 min^32^, even if this estimate ignores cohesin stalling at CTCF sites. Thus, compared to other nuclear motors (*e.g.* RNA Polymerase II^65^ or the replicative helicase CMG^66^ with speeds of 0.02 kb/s and 0.005 kb/s, respectively), and despite its slower rate in cells, cohesin remains a fast motor that extrudes DNA at high rates in the nucleus. This elevated speed likely contributes to the transiency and highly dynamic nature of cohesin-induced long-range interactions.

Fourth, polymer simulations indicate a density of cohesin of 7-12 Mb^-1^ (**Figure 5C**), which for a ‘median’ loop of 230 kb corresponds to 1.6-2.8 cohesin complexes simultaneously bound within the domain. This agrees with previous estimations of 4-8^9,26,57,67^ or 8-32^27^ cohesin complexes per Mb, derived from polymer simulations, absolute quantification of molecules, super-resolution microscopy and live-cell imaging. Our estimates translate into ∼43,000-74,000 cohesin molecules simultaneously bound to chromatin and extruding the 6.2 Gb long human genome during the G1 phase of the diploid HCT116 cells. This range is consistent with absolute quantifications of bound cohesin molecules of ∼60,000-160,000 bound cohesin complexes in HeLa cells in G1^58^ (43,000-114,000 for a ‘diploid’ HeLa cell, instead of the average 64 chromosomes^68^). The estimated densities imply that multiple cohesin complexes are present simultaneously in single TADs, confirming the notion that they emerge from multiple loops extruded simultaneously.

Fifth, while our analyses allow only for a narrow range of cohesin densities, they are consistent with cohesin residence times ranging from 3 min to at least 33 min, in accordance with previous estimations of 3-25 min^27,32,36,58–61^. With the above estimates of cohesin motor speed and density, and taking into account stalling at CTCF sites, we could convert these residence times to genomic processivities and loading rates. Processivities ranged from 150 to 1000 kb (**Figure S10G**), exceeding previous estimations of 120-240 kb^9,26^, while cohesin loading rates fell between 0.2 and 2.8 (Mb x min)^-1^, **Figure S10I**), and are consistent with previous estimates of 0.06-1.2 (Mb x min)^-1^ ^27^. Thus, within the 345 kb long TAD L1, approximately 41-580 cohesin complexes extrude loops during a 10-hour G1 phase.

Sixth, our study analyzed multiple genomic loci featuring different CTCF site distributions, histone marks, Hi-C contact patterns and dynamics with the same methods, allowing us to compare extrusion dynamics and their determinants across chromatin regions. Strikingly, we found that their strong differences can be explained by a single set of values for cohesin density, residence time and motor speed (**Figure 5C**). Our simulations indicate that differences between regions result from the different locations and strengths of CTCF sites rather than from local variations in cohesin dynamics. Thus, our study argues in favor of universal dynamics of cohesin across the genome, rather than for its local tuning. At the same time, these results highlight the crucial and potentially exclusive regulatory role of CTCF in controlling TAD dynamics. Since the sole knowledge of CTCF site location and affinities from ChIP-seq data appears sufficient to quantitatively predict TAD dynamics from polymer physics, our study enables predictive models of loop extrusion, key to nuclear functions such as enhancer-promoter contacts regulating gene transcription^19,20,69^.

Nonetheless, we acknowledge several limitations. First, our chosen loci are particularly strong TADs and loops compared to genomic averages (**Figures S1B-C**). Hence, we expect our quantitative estimations (*e.g.* proximal state fraction, frequency and lifetime) to be larger than for an ‘average’ TAD. Second, the studied domains were all located in the A compartment (**Figure S1A**). Although differences in Hi-C patterns between A and B compartments have been shown to originate from differential CTCF binding and not from differences in cohesin dynamics^47^, it still remains important to determine parameters of cohesin dynamics in repressive chromatin contexts. Third, while we chose to study endogenous TADs with their native distribution of CTCF sites, it would be instructive to systematically vary CTCF binding sites to better understand its major role in shaping TAD dynamics.

Together, our results describe the highly dynamic nature of cohesin-induced interactions in the human genome. They support a model where cohesin complexes almost constantly extrude loops at high rates, and produce transient rather than prolonged contacts between TAD anchors. The uncovered universality of cohesin dynamics and the crucial regulatory role of CTCF will empower predictive models of TAD dynamics and extrusion-dependent genomic functions.

## Acknowledgements

We would like to thank Giacomo Cavalli and Antoine Coulon for useful discussions. We acknowledge and thank members of the MRI imaging facility, part of the national infrastructure France-BioImaging supported by the French Nation Research Agency (ANR-10-INBS-04, Investments for the future). We acknowledge the help of the HPC Core Facility of the Institut Pasteur for the use of computing resources. We thank Xavier Pichon for the initial cloning of the splitGFP array. T.S. was supported by a Contrat Doctoral Spécifique aux Normaliens and Fondation ARC pour la recherche sur le cancer (ARCDOC 42021120004333). We also acknowledge Investissement d’Avenir grant ANR-16-CONV-0005 for funding computing resources used in this work. Some figures were created using images from BioRender.com.

## Author Contributions

T.S. designed the project, performed experiments and polymer simulations and their analysis and wrote the manuscript with input from all authors. M.C.R. participated in cell line construction. B.L. developed parts of the image processing and data analysis pipelines. M.S. performed and generated Capture Micro-C maps. J.Y.T. contributed to image analysis tools. C.Z. and E.B. supervised the project.

## Declaration of interests

The authors declare no competing interests.

## Materials and Methods

### Cell line culture, generation and treatment conditions

#### Cell culture

HCT116 cells were cultured in McCoy’s medium supplemented with GlutaMAX (Thermo Fisher Scientific 36600021), 10% Fetal Bovine Serum (FBS, Sigma-Aldrich F7524) and Penicillin-Streptomycin (50 U/mL and 50 µg/mL respectively, ThermoFisher 15140122). Cells were grown at 37°C in a humidified incubator with 5% CO2 and split every 2-3 days. Cells were tested monthly for the presence of *Mycoplasma* spp., *Ureaplasma* spp. and *A. laidlawii* by qPCR^70^.

#### Cell line generation and genome-editing

CRISPR-based genome editing was performed using a nickase Cas9 to reduce off-target rates. We co-transfected a repair plasmid (0.6 µg), a nickase Cas9 expressing plasmid (0.4 µg, Addgene 42335^71^) and a pair of single guide RNAs (sgRNAs, 0.5 µg each) using JetPrime (Polyplus 101000001) according to the manufacturer’s protocol. The pairs of sgRNAs were designed using ChopChop^72^ with the parameters: ‘hg19’, ‘nickase’ and ‘knock-in’. For repeat array insertion, homology arms were PCR amplified and cloned by a 4-fragment Gibson assembly (NEB E2621S) within the loxP-Blasticidin-HSVTK-loxP-TetOx96 and CuOx150-FRT-Neomycin-FRT plasmids^73^. We used homology arms of 581-1436 bp. 300,000 cells were seeded in a 6-well plate and transfected 24 hours later. Cells were detached from the well and split into four different 10 cm plates for selection less than 24 hours after transfection. Each 10 cm plate contained a different dilution of the initial 6-well plate (from 1/40 to 4/5). Cells were kept under antibiotic selection until unique colonies were seen (about 2 weeks). Single colonies were then picked using cloning disks (Merck Z374431) and put into 24-well plates. Once sufficiently grown, each clone was split in half. One half was used for clone expansion and the other half was seeded on a glass slide for image-based screening. Three days after seeding, Halo tag was labeled, if needed, with 100 nM JFX646^42^ (gift from Luke Lavis lab) in culture medium by incubating the cells for 15 min at 37°C. Cells were then fixed with 4% formaldehyde (Thermo Scientific 28908), diluted in Phosphate Buffered Saline (PBS, Sigma-Aldrich D8537) for 20 min and slides were mounted in Vectashield antifade medium with DAPI (Vector H-1200-10). Clones were imaged with a widefield microscope (Zeiss Axioimager, 63X Plan Apochromat NA=1.4 objective with a LED Xcite 120LED as illumination source and an ORCA-Flash4 LT Hamamatsu camera with 2048x2048 pixels and a pixel size of 6.5 µm). Clones that displayed one or two fluorescent spots per nucleus were further split in a 6-well plate and genomic DNA (gDNA) was extracted (Lucigen QE0905T). PCR (Promega M7801) genotyping was performed by amplifying 5’ and 3’ junctions. The unmodified wild-type band was amplified to assess the zygosity of the insertion. High-quality gDNA of clones verified by microscopy and PCR was purified (Promega A1120) and each genotyping PCR fragment was sequenced. Finally, we checked the integrity of the TetOx96 array by PCR amplifying and sequencing the whole array, using the 3’ junction reverse primer.

In order to enable auxin-dependent RAD21 degradation, we first homozygously inserted the RAD21-mAID-SNAP-IRES-Hygromycin fusion at the endogenous RAD21 locus, using the wild-type Cas9 (0.4 µg Cas9, Addgene 41815^74^, and 0.8 µg each for the repair DNA and sgRNA plasmids), and 100 µg/mL hygromycin B (ThermoFisher 10687010) for selection. Next, we inserted the AtAFB2-weakNLS-IRES-Puromycin at the AAVS1 locus, using a WT Cas9 (Addgene 72833^75^) and by selecting cells with 1 µg/mL puromycin (Invivogen ant-pr-1). The AtAFB2 degron was reported to minimize basal degradation of the degron-tagged protein, as compared to the more common OsTIR1 degron^52^. The weak NLS allowed depletion of both nuclear and newly synthesized cytoplasmic RAD21^52^. Cell lines were regularly cultured with 1 µg/mL puromycin for 1 week to ensure high expression of the AtAFB2 degron^52^ before switching to regular culture medium.

We then expressed the fluorescent reporters needed to visualize the repeat arrays. Using an optimized piggybac transposase (we transfected 0.3 µg of transposase plasmid to insert a single copy of each reporter^76^, among 2 µg of total DNA), we inserted CymR-NLS-2xHalo (Addgene 119907^77^) to visualize CuO repeats and TetR-GFP11x16-GB1-NLS to visualize TetO repeats in the cells. We infected the cells with lentiviruses containing the GB1-GFP1_10-GB1-NLS fragment (bearing the A206K mutation to avoid dimerization^78^) to reconstitute the split GFP. Cells were sorted three times, once a week, to keep only low expressing levels of the reporter proteins. Then, we re-infected the cells with the GFP1_10 lentiviruses to increase the ratio of GFP1_10 over TetR-GFP11x16-GB1-NLS and optimize the signal from the multimerized GFP11 fragments^79^. These cells constituted the parental cell line used for repeat array insertion.

We then sequentially inserted the CuOx150-FRT-Neomycin-FRT and loxP-Blasticidin-HSVTK-loxP-TetOx96 arrays at each anchor of the TADs (**Figure 1A**). Insertion of the TetOx96 and CuOx150 repair plasmids were selected using 6 µg/mL blasticidin (Invivogen ant-bl-1) and 400 µg/mL G418 (Invivogen ant-gn-5), respectively. The expression of antibiotic resistance genes was designed to direct transcription outwards of the TAD interior to avoid interference with loop extrusion. Finally, once TetOx96 and CuOx150 array insertion on the same allele was verified by microscopy, we removed the neomycin and blasticidin antibiotic cassettes using Cre and Flippase (Flp) recombinases. Since transcription is known to alter chromatin dynamics^48,80^ and RNA Polymerase II can slow down, or push extruding cohesin, acting as a mobile barrier for cohesin^49^, the absence of strong transcription from antibiotic cassettes allows to measure cohesin-mediated motion of TAD anchors as purely as possible. 1 µg of Cre (Addgene 123133^81^) and 1 µg of Flippase (Addgene 13793^82^) recombinases were transfected. Clones that lost the antibiotic cassettes were selected by the loss of the herpes virus simplex thymidine kinase (HSVTK) gene, making cells sensitive to 8 µg/mL ganciclovir (Invivogen sud-gcv). 5’, 3’ junctions and the WT allele were sequenced as previously described. These clones were subsequently used for live-cell imaging.

For all cell lines, we obtained heterozygous insertions of the repeat arrays, allowing to track one pair of green and far-red spots in the cells. For the L2 cell line, the CuOx150 array was inserted homozygously, resulting in two distinct far-red spots, while the TetOx96 cassette only integrated in a single allele. Upon sequencing of the L2 clone, we found a plasmid fragment of 396 bp integrated within the inserted exogenous sequence, at the 3’ end of the CuO repeats. This sequence contained a lac operon and promoter, Gateway attB2 sites and a Catabolite Activator Protein binding site. In the T1 cell line, a small fraction of cells retained the antibiotic resistance genes after Cre and Flippase recombinations, as assessed by PCR amplification of the non-recombined alleles.

Repeat arrays were inserted at the following locations in the human genome (hg19): L1 (chr8:60,964,180; chr8:61,310,370), L2 (chr2:235,458,700; chr2:236,026,413), T1 (chr1:36,980,442; chr1:37,901,640), No TAD (chr8:60,733,873; chr8:61,310,370), Adjacent (chr1:37,900,310; chr1:37,901,640).

#### Auxin-mediated RAD21 degradation and western blotting

To deplete RAD21 fused to the mini auxin inducible degron (mAID)^75^, we added auxin (Sigma-Aldrich I3750-5G-A) to a final concentration of 500 µM (from a 500X stock solution diluted in PBS) in fresh culture or imaging medium.

RAD21 depletion kinetics was measured by live-cell imaging and western blotting (**Figure S4**). For western blotting, 300,000 cells were seeded and grown 48 hours in 6-well plates and incubated with auxin for the indicated times. Cells were washed three times with cold PBS and lysed with 200 µL of HNTG buffer (HEPES pH 7.4 50 mM, NaCl 150 mM, Glycerol 10%, Triton-X-100 1%) with 1X protease inhibitor (Roche 5056489001). After cell collection, lysates were rotated for 30 min at 4°C, sonicated and rotated 30 min at 4°C before centrifugation, and supernatants were stored at - 80°C until loading. Protein levels were quantified using the Pierce BCA protein assay (Thermo Fisher Scientific 23225). Samples were boiled for 5 min at 100°C in 1X Laemmli buffer and 10 µg of protein extract were loaded into a 10% Mini-protean TGX gel (Bio-rad 4561036). Samples were run for 90 min at 110 V in Tris-Glycine 1X (Euromedex EU0550) and 0.5% Sodium Dodecyl Sulfate (SDS, Euromedex EU0660s) buffer, and protein was transferred to a nitrocellulose membrane (Pall BioTrace 66485) for 75 min at 100 V in Tris-Glycine 1X and 20% Ethanol (VWR 83804.360) buffer. Membranes were blocked with 5% milk in 1X Tris-Buffered Saline (TBS, Tris 20 mM, NaCl 150 mM) for 1 hour at room temperature (RT). Immunostaining was performed overnight at 4°C with primary antibodies (Rad21 1:1500 (Abcam ab154769), GAPDH 1:50000 (Abcam ab8245)) diluted in 5% milk in 1X TBS-Tween (0.02% Tween, Thermo Scientific 11368311). The membrane was washed three times for 5 min with 1X TBS-Tween at RT and incubated for 1 hour with secondary antibodies diluted in 1X TBS-Tween at RT (anti-rabbit IR800 1:10000 (Advansta R-05060), anti-mouse IR800 1:10000 (Advansta R-05061)). The membrane was washed three times in 1X TBS-Tween for 5 min and once in 1X TBS for 10 min. Before imaging, membranes were soaked in 70% ethanol and air-dried in a dark chamber. We measured fluorescence intensity with the Chemidoc MP Imaging system (Bio-Rad).

For live-cell quantification of auxin-mediated RAD21 degradation kinetics (**Figures S4A** and **S4C**), we used Rad21-mAID-SNAP cells containing the TetR-GFP11x16-NLS and CymR-NLS-2xHalo constructs (parental cell line). 300,000 cells were seeded and cultured in glass-bottom imaging dishes (Ibidi 81158) for two days. Before imaging, the SNAP JF646 dye (gift from Luke Lavis lab) was added to fresh medium at a final concentration of 100 nM and cells were incubated for 90 min at 37°C. Cells were washed three times with warm PBS and imaging medium (DMEMgfp (Evrogen MC102) supplemented with 10% FBS or Fluorobrite DMEM (Thermo Fisher Scientific A1896701) supplemented with 1X Glutamax (Thermo Fisher Scientific 35050061) and 10% FBS) was added to the cells. Time-lapse images were acquired in a bespoke microscope equipped with a 488 nm TA Deepstar Diode Laser (Omicron-Laserage Laserprodukte GmbH) and a 647 nm OBIS LX laser (Coherent Corp.) for excitation. The microscope was equipped with an Olympus UPLAPO 60x 1.42NA objective, and additional optics leading to a 102 nm pixel size in the final image. Green and far-red fluorescence emission was split at 580 nm by a FF580-FDi02-t3-25×36 dichroic mirror (Semrock) and filtered with 525/50 nm and 685/40 nm fluorescence filters (Alluxa Inc.) respectively. Two-color images were captured by two separate sCMOS cameras: a Zyla 4.2 plus for the green channel and a Zyla 4.2 for the far-red (Oxford Instruments) channel. The sample environment (CO2 concentration, temperature and humidity) was controlled with a top-stage chamber (Okolab SRL). All devices of the microscope were controlled using python-microscope^83^ and using cockpit as graphical interface^84^. We took 31 z-slices separated by 0.4 µm each and z-stacks were taken every 15 min for 45 min at multiple positions. Then, cells were removed from the microscope stage and auxin was added to the medium. After placing the cells back in the microscope chamber, we selected new positions on the slide and imaged cells at the same frequency for 4 hours.

To measure RAD21 levels in live-cell images, we used a custom Python script on maximum intensity projected images. We segmented (with Labkit^85^) and tracked (with TrackMate^86^) nuclei using the green channel containing the TetR-splitGFP-NLS signal. We removed all dividing cells and cells at the edge of the image from the analysis. Using these segmentation masks, we measured the median fluorescence intensity in the RAD21 channel. Background intensity was subtracted, and fluorescence intensities were normalized to the first timepoint of imaging without auxin.

#### Cell cycle analysis by Fluorescent Activated Cell Sorting

To assess the fraction of cells in G1 to early S phase and compare it with our image-based assessment of replicated spots, we used Fluorescent Activated Cell Sorting (FACS) with propidium iodide (PI, Sigma-Aldrich P4864) staining of fixed cells (**Figure S3E-G**). 300,000 cells were seeded and grown in 6-well plates and cells were collected 48 hours later. Cells were trypsinized, washed in PBS, and resuspended in 500 µL PBS. 5.5 mL of ice-cold 70% ethanol was added for fixation. Cells were kept at 4°C in 70% ethanol for at least 12 hours until staining. For staining, cells were washed twice in PBS and incubated for 5 min at room temperature with 50 µL of a 50 µg/mL RNAse A solution (Promega A7973). Finally, 400 µL of 50 µg/mL propidium iodide solution was added and cells were incubated for 15 min at room temperature before FACS sorting. We used a Miltenyi MACSQuant Analyzer 10 Flow Cytometer with the 488 nm laser and a 692/75 nm band pass filter. We gated cells based on Forward Scatter vs. Side Scatter, and single cells based on PI height vs PI area. We did not consider polyploid cells (at least triploid) in the analysis (they represented <2.5% of cells). Finally, we analyzed at least 12,000 cells within the final gate of interest.

From the distribution of propidium intensity, the percentage of cells in each cell cycle phase was computed using the Dean-Jett-Fox model^87^ without constraint with the FlowJo^TM^ software.

### Capture Micro-C

Capture Micro-C libraries were generated by merging Micro-C (Dovetail™ 21006) with tiling capture of genomic loci (Agilent Technologies™). Cells from each cell line containing the repeat arrays were aliquoted between 1.2 to 2 x 10⁶ cells in individual Eppendorf tubes. Aliquots were spun down and washed in PBS. The supernatant was carefully removed and discarded. Cell pellets were stored at -80°C for at least a day. Pre-freezing is required to get an optimal MNase digestion profile. Cell pellets were thawed at room temperature and then processed as prescribed by the Micro-C protocol. For the end repair steps, we used the NEBNext Ultra II DNA Library Prep Kit for Illumina (NEB E7645) instead of the Dovetail kit. After adapter ligation with the NEB kit, DNA was purified *via* SPRI beads (Beckman A63880) as described in the Dovetail kit. Ligation capture and library amplification was performed with reagents from the Dovetail kit, except for the sequence index which was taken from the NEB kit. After verifying that libraries had a correct concentration and size distribution, we continued with the Agilent SureSelectXT HS2 Kit (G9987A, design #S3442002). Capture probe design was performed by Agilent. The coordinates of capture probes were: chr8:60,458,500-61,587,500 for the L1 and No TAD regions, chr2:235,182,500-236,297,500 for L2 and chr1:36,700,000-38,175,000 for T1. We followed the manufacturer’s protocol for pre-pooling 8 sequencing libraries. Finally, we checked concentrations and size distributions before sending capture-sequencing libraries for sequencing. We sequenced with BGI Illumina 100-bp paired-end sequencing (PE100).

The data generated in this manuscript were pooled from two biological replicates for each cell line, except for the No TAD cell line where a single experiment was performed. Raw sequencing data from BGI was checked by FastQC^88^ (FastQC v0.12.1). None of the replicates showed any irregularities. All sequencing samples were “hard trimmed” to a 50 bp length via Trim Galore (Cutadapt version 0.6.10^89^). Next, valid Micro-C contacts were obtained with the HiC-Pro pipeline^90^ (HiC-Pro_v3.1.0). HiC-Pro uses Bowtie2^91^ to map the reads to the chosen genome. All valid Micro-C contact pairs obtained from HiC-Pro were filtered for the corresponding region of interest and then processed via the Cooler package v0.9.1^92,93^.

### Live-cell imaging of TAD anchors

For live-cell microscopy of TAD anchors, cells were plated on a 35 mm glass-bottom imaging dish (Fluorodish FD35-100). 48 to 72 hours after seeding, the medium was replaced with fresh medium containing 100 nM of JFX646 Halo dye and the cells were incubated for 15 min at 37°C. Cells were washed twice with PBS and the medium was replaced with live-cell imaging medium (DMEMgfp supplemented with 10% FBS or Fluorobrite DMEM supplemented with 1X Glutamax and 10% FBS). For RAD21 degradation, cells were treated with 500 µM auxin. Auxin was maintained in the Halo labeling medium and in the imaging medium during acquisition. Cells were imaged for 2 hours, starting 2 hours after auxin addition. Before imaging, cells were allowed to equilibrate in the microscopy incubation chamber for at least 15 min at 37°C and 5% CO2.

Time lapse image acquisition was performed with an inverted microscope (Nikon) coupled to the Dragonfly spinning disk (Andor) using a 100X Plan Apo 1.45 NA oil immersion objective. Excitation sources were 488 nm (150 mW) and 637 nm (140 mW) lasers. Exposure time was set to 85 ms for both channels with 1% laser power in far-red and 5-8% laser power in the GFP channel depending on the imaged cell line. Z-stacks of 29 optical slices separated by 0.29 µm each were acquired every 30 s using the perfect focus system and five different stage positions were imaged for each 2-hour acquisition. The two channels were acquired simultaneously on two distinct EMCCD iXon888 cameras (1024 x 1024 pixels, effective pixel size: 0.121 µm).

### 3D polymer simulations of loop extrusion

#### Molecular dynamics simulations

Polymer motion was simulated with Langevin dynamics in LAMMPS^94^. The polymer was modelled as a freely jointed chain and consecutive monomers were connected by a harmonic bond with a potential *E*_*bond*_ = 30(*r* − 1)^2^, where *r* is the distance between bead centers. We converted simulation units into physical units by comparing the plateau of simulated and experimental 2-point MSD curves without loop extrusion (auxin-treated cells for experimental conditions) for each genomic locus. First, we set the spatial conversion by comparing the simulated and experimental 2-point MSD plateau, using a bead size of 2 kb. This yielded a bead physical size of 45 nm and, thus, a compaction of 44 bp/nm consistent with previous estimates of 18-66 bp/nm^95–97^. Second, we set the temporal conversion by comparing the simulated and experimental timepoint at which half the 2-point MSD plateau was reached. This led to the conversion that 2000 simulation timesteps = 3 s. We simulated a polymer of 1300 beads, representing 2.6 Mb of DNA centered on each TAD. We used fixed boundary conditions and used a confinement sphere of radius 24 bead diameters with 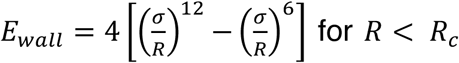, where *R* is the distance between the confining sphere and the center of a bead, *σ* is a size factor set to 0.5 bead diameter and *R*_*c*_ is the cutoff distance set to 0.5 bead diameter. This led to a volume occupancy ratio of 10% in agreement with the experimentally measured chromatin density of 10-15% in the nucleus^98–100^. Simulations were run for a total of 15x10^6^ simulation timesteps representing 11 hours to 5 days of computational time.

The polymer was first equilibrated for 10^6^ simulations steps, after which its radius of gyration and end-to-end distance were stabilized. Then, we ran 400 extrusion steps at 27 kb/s and 400 extrusion steps at 5 kb/s to renew the pool of already loaded cohesin complexes. Finally, 1300 extrusion steps were performed at the defined motor speed to equilibrate the polymer under active extrusion before recording conformation snapshots during 4.8x10^6^ simulation steps. We generated 50 independent simulations for each set of extrusion parameters, each spanning two hours, as in experimental data. For the sets of parameters used in **Figure 6**, we generated 600 independent simulations for each condition.

#### Loop extrusion modeling

Loop extrusion was modelled by the creation and destruction of sliding links between non-consecutive beads. In absence of obstacles, loop extrusion occurred bi-directionally. It continued uni-directionally if the cohesin was blocked by a CTCF site on one side^13^. Cohesin complexes were loaded at random positions on the polymer and at a rate defined by the cohesin density divided by its residence time, therefore ensuring constant cohesin density. Cohesin complexes detached at a rate defined by the cohesin residence time, which was assumed to follow an exponential distribution. We allowed cohesin complexes to traverse each other^101–103^ since there is currently more experimental evidence supporting that SMC complexes can traverse each other^101–103^. Nevertheless, we cannot rule out a model of stalling upon cohesin collisions before resuming of extrusion or unbinding from chromatin, as observed with other roadblocks *in vitro*^63^.

Cohesin motor speed was modified by changing the number of time steps between the creation of a new sliding bond. Upon encounter with a convergently oriented CTCF site, half of extruding complexes were stopped and half proceeded unimpeded, recapitulating the estimated 50% occupancy of CTCF sites by the CTCF protein^57^. Cohesin complexes were stalled at CTCF sites for a duration drawn from an exponential law with a mean CTCF residence time that was defined specifically for each CTCF site based on CTCF ChIP-Seq, as described below. The extremities of the polymer exhibited an infinite CTCF residence time and a 50% probability of stalling cohesin. This allowed to correct for cohesin complexes coming from outside of the simulated polymer, which were not modelled. The indexes of beads linked at each timepoint in these 1D simulations, starting at steady-state, were used as input for Langevin dynamics to generate 3D polymer molecular dynamics simulations.

#### Modeling CTCF residence on chromatin

We assigned a specific residence time to each CTCF site based on the corresponding ChIP-Seq peak. We defined a 2.6 Mb (length of the simulated polymer) region centered on the TAD of each genomic region. We retrieved CTCF sites mapped on CTCF ChIP-Seq peaks. For each CTCF site, we used the CTCF peak fold enrichment computed by MACS2, which takes into account local background, as a measure of CTCF residence time. We computed the median fold enrichment for all peaks identified across the genome and normalized the CTCF fold-enrichment present within the studied genomic region to the genome-wide median. We thus assumed that the affinity of the binding site linearly scaled with Chip-Seq fold-enrichment, as predicted in sequence to affinity models^104^. We then used previous estimations of CTCF residence time of 1-4 min in mESCs^32^ to convert the genome-wide median ChIP-Seq fold enrichment to a CTCF median residence time of 2.5 min. Therefore, CTCF sites above the genome-wide median fold enrichment had residence times longer than 2.5 min (**Figure 4B**). Because CTCF sites are defined as 20 bp-motifs, multiple binding sites and ChIP-Seq peaks could be found within a single polymer bead of size 2 kb. If the CTCF sites were oriented in the same orientation, the peak fold enrichments were added. In the few case where one CTCF ChIP-Seq peak overlapped two binding sites of opposite directions, the CTCF residence time associated with each orientation was adjusted based on the relative p-value of the two CTCF sites.

We chose to model the fold enrichment of ChIP-Seq peaks by changing CTCF residence time instead of CTCF occupancy. While bulk ChIP-Seq cannot distinguish the contribution of the association rate kon and the dissociation rate koff to the detected peaks, it was found that changes in binding site sequence had a higher impact on koff than kon105. Thus, we modelled changes in ChIP-Seq peaks by changes in CTCF residence time, while fixing the occupancy.

#### Genomic data analysis

The hg19 genome was used for all genomic analyses. HCT116 genomic data were retrieved from *Rao et al*^6^. Loops and TADs were called using Juicer 1.19.02^106^ HiCCUPS and Arrowhead, respectively on Hi-C maps of *Rao et al*^6^. The following flags were used for HiCCUPS: -r5000, 0000 -k KR -f 0.1 -p 4,2 -I 7,5 -t 0.02,1.5,1.75,2 -d 20000, 20000; and for Arrowhead: -m 2000 -r 5000 -k KR --threads 10.

For ChIP-Seq data of CTCF, SMC1, RAD21, we used publicly available data from *Rao et al*^6^. Raw reads were quality-checked using FastQC^88^. Reads from different replicates were first mapped independently using Bowtie2 v2.2.6.2^91^ with default parameters, and the correlation between replicates was computed using wigCorrelate^107^. Replicates with correlations larger than 0.9 were pooled together and mapped again. We removed blacklisted regions^108^ and called peaks using default parameters of MACS2 v2.1^109^. For histone marks ChIP-Seq, we used the flag ‘broad’. CTCF motifs were identified genome-wide using FIMO^110^ with the flags -max-stored-scores 50000000 and -thresh 0.001 and the Jaspar motif MA0139.1. We then mapped CTCF sites identified with a P-value < 1x10^-5^ onto CTCF ChIP-Seq peaks.

ChIP-Seq peaks of CTCF, SMC1, RAD21 were intersected within regions of 20 kb centered around TAD anchors using pgltools intersect1D^111^.

A and B compartments were identified using ‘eigenvector’ from Juicer with the flags KR BP 100000. A compartments were defined as high H3K27ac, Pro-Seq, H3K36me3, H3K4me3 and H3K4me1 signals and B compartments as the opposite.

#### Choice of genomic loci for labeling of TAD anchors

Using Hi-C, ChIP-Seq and PRO-seq data from Rao *et al*^6^, we filtered loops and TADs based on the following criteria: (i) size comprised between 300 kb and 1.5 Mb, (ii) at least one peak of SMC1, RAD21 and CTCF at both anchors, (iii) at least one pair of convergent CTCF sites at anchors, (iv) no gene at anchors and (v) low gene expression within the domain (<1.5 reads per kilobase million, RPKM). These criteria were defined to minimize possible cohesin-independent chromatin interactions that might obscure cohesin-dependent extrusion dynamics, which we aim to measure as purely as possible. From this subset, to further ensure that selected domains were cohesin-dependent, we removed domains exhibiting enhancers at their anchors (as identified in the genehancer double elite set^112^. After filtering, we obtained a list of 96 loops and 32 TADs, from which we removed manually highly nested domains and domains containing alignment artifacts in Hi-C maps. We then chose strong loops and TADs, exhibiting highly ranked sgRNAs (from ChopChop^72^) to facilitate genome editing and at less than 5 kb from CTCF anchors to ensure the most accurate readout of anchor-anchor distances, as previously assessed by polymer simulations^54,55^.

For the adjacent control cell line, we inserted the TetOx96 repeats 6 kb away (mid-array distance) from the CuOx150 repeat array used to label the 3’ anchor of the T1 locus (**Figure S1A**). For the No TAD control, we inserted the TetOx96 repeats on the 5’ side of the 5’ anchor of the L1 locus in a cell line that already contained the CuOx150 repeat array at the 3’ anchor. No CTCF site was present within 100 kb and no convergent CTCF site relative to 3’ anchor was identified within 117 kb of the TetOx96 repeat array insertion site (**Figure S1A**).

#### Image analysis

##### Image shift correction

Due to misalignment of the motorized microscope stage device, a small shift can appear between consecutive 3D images after displacement at multiple fields of views. While axial shift can be ignored thanks to the perfect focus system, lateral shifts can reach a few hundred nanometers, limiting the efficiency of our tracking analysis. To attenuate lateral shifts, we first computed 2D cross-correlations with the first imaged timepoint using 2D projections of the far-red channel. We then used the estimated displacement to correct both the green and the far-red channels with a pixelic precision to avoid pixel interpolation, which could have negative impact on spot localization by maximum likelihood.

##### Eliminating replicated spots

We manually eliminated cells dividing during the duration of time lapses. To eliminate cells containing replicated spots (*i.e.* spots in cells that, at least, started their S phase), we computed the elongation of the detected spots and removed cells with a large spot elongation (**Figure S3E**). A small round spot likely corresponds to a single chromatin locus whereas an elongated spot likely corresponds to two overlapping spots coming from two replicated chromatids. Manual inspection of maximum intensity projection of images allowed us to eliminate simple cases where two distinct spots above the diffraction limit are seen. However, the two replicated spots may not always be resolved as distinct spots due to the diffraction limit. Spot elongation was measured in two steps. First, a 3D Laplacian of gaussian filter was applied to the 3D image stacks, and spots were detected by searching for local maxima above a given threshold. Next, we fitted a 3D second order polynomial function (paraboloid) centered on each local maximum. We measured the amplitude of the paraboloid at 8 positions regularly positioned on a circle around the vertex of the paraboloid and estimated the covariance among these 8 positions. We defined a spot elongation score as the ratio between the maximum and the minimum of the eigen values minus 1. A score close to 0 corresponds to an isotropic spot, whereas a large score corresponds to an elongated spot. We computed this score for each spot in the far-red channel and superimposed these scores to a 2D time- and z-projected image of time lapses (**Figure S3E**). This image was generated for each time lapse as a guide to detect replicated spots. The spot elongation score allowed us to remove cells containing replicated spots in more complex cases where the two replicated spots could not be distinguished. Our method to eliminate replicated spots was conservative, as the fraction of cells in imaging data thus retained was slightly lower than the fraction of G1 cells measured by FACS (**Figures S3F-G**). We note that we cannot eliminate cells that started the S phase but where the labelled locus was not replicated yet. We consistently observed a lower fraction of cells in G1 in auxin-treated cells as compared to untreated cells, as expected from cell cycle arrest due to RAD21 depletion^113^.

##### Detecting and tracking fluorescent spots

The 3D image time-series were processed using Fiji plugins as follows. First, we manually defined rectangular Region Of Interests (ROIs) around each pair of spots corresponding to cells in G1 or early S phase. A single ROI was used for each pair of spot, *i.e.* each ROI contained all positions explored by the locus during the time lapse acquisition.

Detection and tracking of fluorescent spots were performed separately for each color channel using TrackMate^86^ (**Figure S3A**). To improve the detection of spots in images of varying Signal-to-Noise Ratio (SNR) relative to the default Laplacian of Gaussian detector, we implemented a spot detector based on the determinant of the local Hessian matrix. The resulting Hessian detector is more robust to spurious local intensity maxima that tend to occur at the edge of bright objects^114^, such as nuclei. We used a different filter size in Z and XY, to account for the axial elongation of the PSF^115^. In order to facilitate removal of spurious detections, the ‘Quality’ metric computed by TrackMate was normalized within the ROI. Detections with quality values below 0.7 and 0.8 for Halo and GFP, respectively, were rejected. We then used a localization algorithm based on parabolic interpolation^116^ to refine the spot position with sub-pixel accuracy.

Next, we connected detected spots over time to generate trajectories. This was done using the simple Linear Assignment Problem (LAP) tracker^117^ in TrackMate with the following parameters: linking distance = 0.9 µm, gap-closing distance = 1.4 µm and maximum frame-gap = 12 frames. Spurious detections due to noise typically generated short trajectories. To remove them, we set the minimum number of detections per trajectory to ∼20 for the GFP channel and ∼40 for the Halo channel. The resulting trajectories possibly contained large gaps, *i.e.* several consecutive time points with detections below the above-mentioned quality threshold. To address this, we implemented a gap-filling step in TrackMate, where trajectories with gaps were automatically revisited and corrected as follows. For each gap, we first used linear interpolation between the previously detected spot positions before and after the gap. Second, we performed another detection with the Hessian detector within the gap interval, but restricted to a distance of 0.5 µm at most around these interpolated positions. We then added the detection (if any) with the highest quality above the quality threshold to the trajectory and updated the interpolated positions for the gaps in the subsequent time-points. This led to a partial closure of the gaps, but smaller gaps typically remained because no local maximum of sufficient quality was found within the search space. The detection and tracking parameters were optimized automatically with the TrackMate-Helper plugin^118^ using manually annotated images as ground truths. Parameters were optimized to maximize the ‘matching score against ground truth trajectories, penalizing spurious tracks’, a quality metric previously defined in the single particle tracking challenge^119^. Before pairing of the two channels, the coordinates of trajectories were corrected for chromatic aberrations (see below and **Figures S3A-B**).

Green and far-red spot 3D trajectories were paired together using a custom-written Fiji plugin “Pair TrackMate files” as follows. For each green spot trajectory, we considered all far-red spot trajectories that overlapped in time and counted the number of time points for which the detected green and far-red spots were within 2 µm of each other. The trajectory pair with the largest number of such time points was retained and both trajectories were removed from the list of candidate trajectories. This procedure was iterated until one of the lists of green or far-red trajectories was empty. The coordinates of each spot in the paired trajectories were then refined by maximum likelihood (see below).

All tools described here are publicly available in Fiji^120^, either in the TrackMate plugin, or in two extensions “TrackMate-Helper” and “TrackMate-Pairing” available by subscribing to the Fiji update sites of the same name.

##### Correcting for chromatic aberrations

Correcting chromatic aberrations is crucial to precisely compute the 3D distance between 2-color loci. To estimate chromatic aberrations in 3D in the exact same plate and medium used for imaging the TAD anchors, we acquired 3D reference images of actin in WT HCT116 cells in the green and far-red channels using CellMask green and Deep Red actin stains (ThermoFisher A57245 and A57243, respectively; **Figure S3A**). At least ten fields of view of actin z-stacks were acquired every two days of imaging. Chromatic shifts were then measured with Chromagnon^121^ using the averaged actin images as references and the option ‘Local align’ set to ‘None’. The estimated 3D XYZ translations, 3D magnifications and 2D lateral rotations were used to correct the coordinates of spot localizations (**Figure S3B**). Timelapse images of fluorescent beads (Tetraspeck 0.1 µm, T7279) in the green and far-red channels were acquired using the same imaging parameters as live-cell imaging time lapses to evaluate chromatic aberration correction. Beads were positioned on the bottom of the imaging plate in PBS. Bead localizations were corrected for chromatic aberrations using actin images of cells as previously described and the distance between the channels of each fluorescent bead was computed (**Figures S3A** and **S3B**). We found that we could accurately correct for chromatic aberration since the median distance decreased from 265 to 50 nm after correction (**Figure S3B**). This highlights the necessity to correct for chromatic aberrations to study sub-micrometric distances between chromatin loci using different color channels.

##### Refining localizations and measuring localization precision

All paired positions obtained using the above methods, including interpolated missing positions, were refined using maximum likelihood estimation (MLE), which is known to be optimal for precise localization when the PSF and noise models are known. Because our MLE algorithm assumes Poisson noise, we first converted the pixel values into photon counts using an affine function. For this purpose, we acquired 100 images of the same field of view of fixed cells, measured the mean and variance of each pixel, and determined the affine function that makes the mean approximately equal to the variance. This calibration was necessary because our EMCCD cameras do not provide photon counts. All spots were localized by MLE using an anisotropic 3D Gaussian PSF. The iterative MLE algorithm was initialized using the spot positions identified by TrackMate or by interpolation as described above, and was performed in a 3D region of 7x7x7 pixels centered on this position (**Figure S3A**). To determine the standard deviation of the Gaussian, we first performed the MLE fit by estimating the spot coordinates, amplitude, background together with the lateral and axial standard deviations. In a second step we fixed the three (X, Y and Z) standard deviations to the medians of the estimated values on each cell line (grouping untreated and auxin-treated cells together) and performed MLE again by estimating the coordinates, amplitude and background with the standard deviations held constant. Furthermore, for each coordinate *v* ∈ {*x*, *y*, *z* }, we computed the Cramér-Rao bound^50^, providing us an estimation 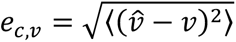 of the precision in nanometer for each fluorescent spot and each channel *c* ∈ {*r*, *g* } at each time point (**Figure S3D**). The 3D localization precision was then computed as 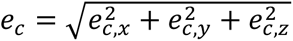 and the precision on distances between the far-red and green loci was computed as 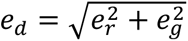. Localization precision was included in all downstream analyses, as a measure of the confidence of each localization.

##### Trajectory quality filtering

Before analysis, time series were quality filtered to minimize localization and tracking errors (**Figures S3A** and **S3C**). First, we removed low precision localizations in each channel (*e*_*c*_ > 250 nm), pairs of localizations with low distance precision (*e*_*d*_ > 250 nm), localizations at the edge of the volume used for Gaussian fitting (< 1 pixel from volume edge). To filter out stepwise tracking errors, we computed the frame-to-frame step size of each fluorescent spot and the frame-to-frame step in distance between the two spots. Timepoints exhibiting a z-score > 1.75 for the step in distance and at least one of the two step sizes simultaneously were removed as tracking errors. We trimmed the end of time series before they exhibited more than 10 consecutive missing values. After removal of single timepoints, we filtered out short and low-quality time series by removing: (i) time series with less than 20 timepoints, (ii) time series with a median precision on distance *e*_*d*_ > 150 nm, (iii) time series with more than 30% of missing frames. Finally, we filtered out trajectory pairing errors by removing time series with a median distance between the far-red and green spots larger than 1.5 µm (except for the adjacent cell line where we used a threshold of 1 µm).

Finally, remaining gaps in time series were filled by interpolating the distance between the time points immediately before and after the gaps. These interpolated distances represented on average 8.6% of all timepoints (6-13% is the minimum to maximum among all cell lines and treatments).

##### Scoring of localization precision

We assigned a weight to each computed anchor-anchor distance based on the localization precision *e*_*d*_ as follows:

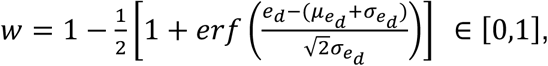

where *γ*_*e*_*d* and *σ*_*e*_*d* stand for the mean and standard deviation, respectively, of *e*_*d*_ computed across all cell lines and treatments. Better localization precisions of far-red and green spots lead to higher scores, and distances measured with better precision thus contribute more to subsequent analyses than less precise distance measurements.

##### Visualization of images

Solely for visualization purposes (Figures **1D-E**, **S2**, **S3A**, **S3E** and **S4A**, **Video S1**), photobleaching of time lapses was corrected by the exponential fitting function of ImageJ ‘Bleach Correction’ plugin.

#### Quantification of distance time series

##### 2-point Mean-Squared Displacement

The 2-point MSD at a time lag *k*δt (where *k* is an integer and δt = 30s is the sampling time interval) for a single trajectory with *N* time points was computed as:

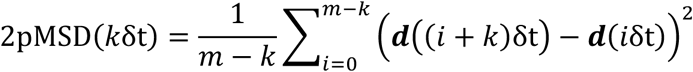

where ***d***(*t*) = ***r***_red_(*t*) − ***r***_green_(*t*) is the 3D vector linking the positions of the far-red and green spots at time *t* and *m* − *k* is the number of time intervals of length *k*δt contained within the time series. For a set of *n* > 1 trajectory, the average 2-point MSD is:

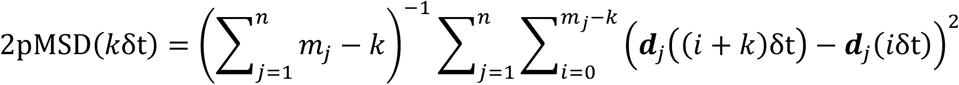

Where ***d***_j_(*t*) is the 3D vector linking the positions of the far-red and green spots at time *t* in trajectory *j* ∈ [1, *n*] and *m*_j_ is the number of timepoints in trajectory *j*.

##### Proximal state segmentation

To segment distance time series into intervals of proximal states, we used a simple, model-free approach involving a spatial threshold and a temporal threshold, which we previously validated on polymer simulations^54^. The spatial threshold was defined as the 95% quantile of the theoretical distribution of distances in the closed loop state. We built this theoretical closed state distribution by simulating N=10^7^ pairs of 3D spot positions and sequentially adding three sources of errors as follows. First, We simulated the 3D positions of the two anchors assuming that they are separated by a distance of 40 nm, corresponding to the cohesin ring size^122,123^ and its expected nontopological entrapment of DNA^124,125^ and assuming random 3D orientations. ring size^122,123^ and its expected nontopological entrapment of DNA within its ring^124,125^. Second, to account for the chromatin linker between the anchors and the fluorescent reporter sequence, we shifted each position with a random value following a normal distribution. This value was determined based on the known genomic distance between the anchor and the reporter and assuming a Kuhn length of 40 nm (in agreement with experimental estimations in yeast and *Drosophila* of 16-134 nm^95–97,126^) and a chromatin compaction of 44 bp/nm (in agreement with estimations of 18-66 bp/nm^95–97)^. Third, to account for random localization errors, we added normally distributed 3D random displacements with lateral and axial standard deviations randomly drawn from the n=12,269-93,431 localization precisions estimated by Cramér-Rao bounds on each cell line. Thus, our theoretical distribution of distances in the closed state takes into account the observed small differences of localization errors (**Figure S3D** and **Table S1**) and reporter-anchor separations (**Table S2**) between the different cell lines. We used spatial thresholds of 0.199, 0.218, 0.236 and 0.220 µm for L1, L2, T1 and No TAD, respectively (**Table S2**).

We used a temporal threshold of 3 min for all cell lines. We also varied this threshold to estimate the range of possible inferred proximal state fractions, frequencies and lifetimes (**Figures S7B-E**). To segment proximal states, we identified all time intervals with anchor-anchor distances below the spatial threshold and durations exceeding the temporal threshold (ignoring intervals consisting of a single timepoint). For each distance vs time series, this yielded a binary time series, where 1 indicates a proximal state and 0 no proximal state. We then applied a rolling average over sliding windows of length equaled to the temporal threshold to filter out the effect of brief distance fluctuations (false negative detections) and avoid fragmenting proximal states, as previously validated on polymer simulations^54^. Finally, we labelled as proximal states timepoints with a resulting value above 0.5 (and non-proximal otherwise; **Figure S6**). Obviously, this method cannot detect proximal states shorter than the temporal threshold (**Figure S7A**).

##### Computing fraction, frequency and lifetime of proximal states

The fraction of proximal states reported in **Figure 2B** was simply computed as: 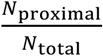, where *N*_proximal_ denotes the number of timepoints in the proximal state and *N*total denotes the total number of timepoints.

The frequency of proximal states reported in **Figure 2C** was computed as: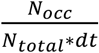, where *N*_*occ*_ is the number of proximal state intervals and *dt* is the time interval between two acquired 3D image stacks (0.5 min).

The mean lifetime of proximal states reported in **Figure 2D** was computed by fitting an exponential function to the histogram of proximal state durations, taking into account censoring as in *Gabriele et al*^26^.

##### Estimation of loop state fractions

To estimate loop state fractions, we used a method previously described and validated on polymer simulations in which an analytical model is fitted to the distribution of coordinate differences^54^ (**Figure S8A**). Briefly, we assumed that the coordinate differences between TAD anchors *δv* (*v* ∈ {*x*, *y*, *z*}) follow a normal distribution of mean 0 and variance 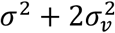, where *σ*_*v*_ is the localization precision for dimension *v* and *σ* depends on the loop state as indicated below:

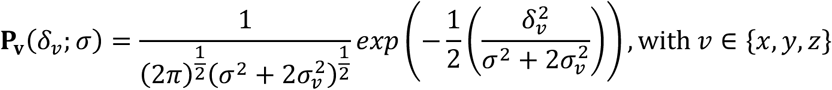

In the closed and open states, *σ* is assumed to be constant and called 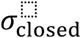 and 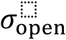, respectively. For each cell line, we obtained the variances of coordinate differences 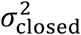 and 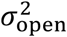 by fitting this Gaussian model to the distribution of coordinate differences in the segmented proximal states and auxin-treated cells, respectively. For each distribution, the localization precision for dimension *v* was set to the averaged localization precision:

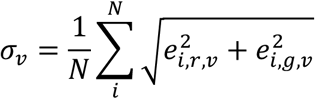

where *N* is the number of coordinate differences in the distribution, *e*_*i*_ is the estimated localization error for spot *i* (see section ‘Refining localization with maximum likelihood and measuring localization precision’). *r* and *g* indicate the color channel of the fluorescent spot.

The distribution of coordinate differences in the extruding state is modelled as an integral over *σ*^2^ varying from 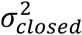 to 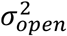. This model assumes that the polymer is at equilibrium at each step of the extrusion process and that the anchors in the extruding state behave as if part of a shorter polymer in which the loop is absent. The full analytical model reads:

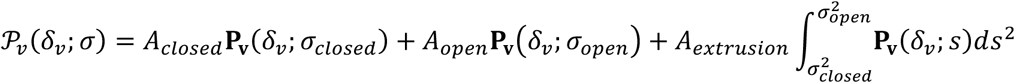

where *A*_*closed*_, *A*_*extrudin*g_, *A*_*open*_ are the three loop state fractions to be estimated. This model was fitted to the three distributions (x, y and z) of coordinate differences simultaneously.

We used bootstrapping to estimate ranges of the inferred loop state fractions. To do so, we randomly drew time series (and not single timepoints) with replacement from the original dataset, using 100% of the time series. For the proximal and auxin-treated cells, we created two datasets. The first set was used to estimate the variance of the coordinate difference distribution for the proximal and open states, while the second set was used to infer loop states. For untreated cells, we created a single dataset from which the fractions of loop states were inferred.

##### Estimation of closing rate

To estimate the closing rate, we adapted a method previously validated on polymer simulations^54^. We aligned time series such that the segmented proximal states coincide at *t* = 0 and computed the ensemble mean squared anchor-anchor distance (EMSAAD) 〈〈*R*^2^〉〉(*t*) from these aligned time series.

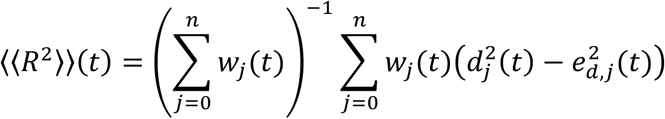

where 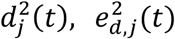, and *w*_j_(*t*) are the squared distance, the squared localization precision and the weight for the track *j* ∈ {0,1, … , *n*} at time *t*, respectively (see above “Refining localizations with MLE and measuring localization precision” and “Scoring of localization precision”). This weighting allows us to reduce the influence of distances associated with low precisions *e*_*d*_ on the closing rate estimation. We also computed the weighted standard error of the EMSAAD:

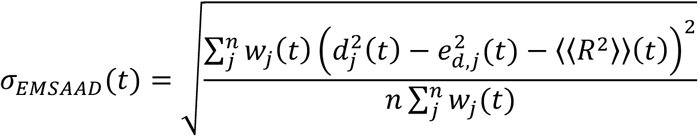

We fitted the EMSAAD to two distinct models, (i) a constant model *f*_1_:

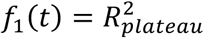

(ii) a piecewise linear curve *f*_2_ defined by three parameters 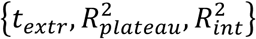:

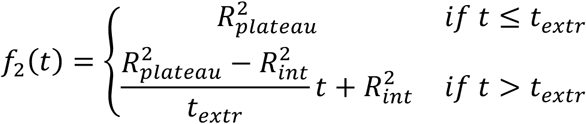

where 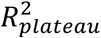 is the averaged mean squared distance at equilibrium, *i.e.* before we can detect the influence of incoming cohesin(s) on anchor-anchor distances, *t*_*extr*_ is the time at which distances start to decrease due to the action of cohesin complexes involved in the shortest 1D path between anchors, and 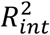 is the squared distance at time *t* = 0 (**Figure 3C**). We fitted the experimental and simulated EMSAAD to the two models *f*_1_ and *f*_2_ by minimizing the mean squared error 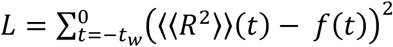 over the time interval [−*t*_w_, 0] preceding the proximal states (**Figures 3D** and **6A**). We then computed the Bayesian information criterion^56^ to select the best model among *f*_1_ and *f*_2_:

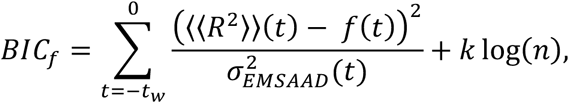

Where *k* = {1,3} is the number of parameters fitted for the models *f*_1_ and *f*_2_, respectively. We chose the closing rate V_0_ that gave the lowest BIC (**Figures S9A** and **S9B**):

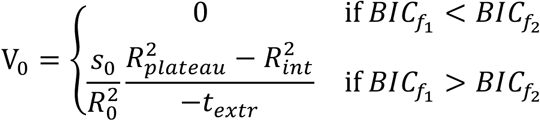

where *s*_0_ is the genomic size between anchors, and 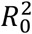 is the mean squared distance measured in auxin-treated cells. An advantage of the 3-parameter model above is that it automatically determines the averaged mean squared distance at equilibrium 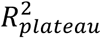 and the starting point *t*_*extr*_ at which the effective genomic separation between anchors decreases.

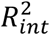 was initialized at the last value of EMSAAD, just before *t* = 0, while 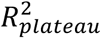 was initialized at the mean value of EMSAAD from *t*_w_ to *t*_*extr*_. To avoid local minima, we initialized *t*_*extr*_ at multiple positions along the aligned time series and kept only the parameters resulting from the fit with the lowest log-likelihood. We used a temporal window of *t*_w_ = 50 time points, corresponding to 25 min. In order to avoid considering time points where the polymer is in a proximal state or in the process of relaxing from a previously formed loop after cohesin unloading or CTCF site bypassing, we ignored proximal states if an earlier proximal state occurred within this time window. This filtering and the size of the fitting window did not affect the estimation of closing rate, as long as a sufficiently large window size was used to observe a plateau in squared distances (**Figure S9D**). Also, we did not consider the two timepoints preceding the proximal state for fitting since they are directly affected by the thresholding procedure used for proximal state segmentation. The fitting was performed only in the cases where at least 20 time series could be aligned. This was generally the case, except for experimental randomized No TAD time series, where only 569/5000 bootstraps fulfilled this requirement.

As negative controls for the closing rate estimation, we randomly shuffled time points within individual time series (without grouping measurements from different time series). We then segmented proximal states and estimated the closing rate using the exact same procedure as for the original (unshuffled) data. Time series were randomized at each bootstrapped sample.

##### Comparison of simulations to experiments

For each of the experimentally studied genomic region, we explored a simulation parameter space where the cohesin density ranged from 1 to 40 Mb^-1^, the cohesin residence from 2 to 33 min and with motor speeds of 0.25 and 1 kb/s (**Figure 4A**). At a cohesin motor speed of 1 kb/s, we explored the parameter space with a finer resolution. These ranges embraced current experimental and computational estimations of these parameters of 4-32 cohesin per Mb^9,26,27,57^, residence times of 3-25 min^27,32,36,58–61^, and the 0.5-1 kb/s motor speeds estimated *in vitro*^10,11^, in addition to the theoretical rough approximates of 0.2-1.3 kb/s obtained from dividing the cohesin residence time by the median loop length of 230 kb (**Figure S1B**). We did not explore cohesin residence times higher than 33 min due to the finite size of the simulated polymer. The corresponding processivities (∼1000 kb at 1 kb/s, **Figure S10G**) were slightly lower than the length of the simulated polymer (2600 kb) and higher cohesin residence times could be affected by edge effects.

From the 3D simulations, we computed the distance between beads located near, but not at, the simulated TAD anchor, consistent with the genomic distance separating the center of repeat arrays and the CTCF site defining TAD anchor for each studied locus (3.9 to 8 kb depending on the genomic region, **Table S2**). To model non-uniform localization precision due to fluorophore photobleaching, we fitted a linear law to the progressive decrease of mean experimental localization precision 〈*σ*_*c*,*v*_(*t*)〉 in x, y and z as function of time, independently for each color channel, and for each cell line. We then used this linear function to add localization errors to the simulated exact coordinates. Finally, we down-sampled simulations to 1 frame per 30 seconds and 241 frames in total as in the experimental live-cell imaging. The spatial and temporal thresholds used to segment proximal states in experimental tracking data were used to segment proximal states in simulated time series. Unless otherwise stated, these simulations containing noise from localization errors, finite genomic length between CTCF sites and fluorescent reporters and at a temporal resolution of 30 s per frame were used.

To build contact maps from polymer simulations, we sampled 100 conformations from each of the 50 independent simulations generated for each set of parameters. We used a capture radius of 1 bead (*i.e.* 45 nm) to model Micro-C maps and define contacts in simulated contact maps. This threshold is below the 100-150 nm capture radius estimated for Hi-C from comparison of distance maps to genomic contact maps^29,30^. Indeed Micro-C is expected to display a lower capture radius than Hi-C^5,44^. We restricted our analysis to contact maps spanning 1.1 times the length of the studied TAD.

To compare polymer simulations with experiments, we considered: anchor-anchor distance distributions, the p(s) curve from contact maps, 2-point MSD curves; proximal state fraction and frequency distributions obtained from bootstrapping, and proximal state lifetime. Note that simulated and experimental p(s) were normalized to 1 at 20 kb (**Figure S11C**). Likewise, for the 2-point MSD curves we set the first time lag value to 1 and considered the values for a maximum time lag of 300 seconds (**Figure S11D**).

For each quantity and set of simulated parameters, the deviation of polymer simulations from the experimental data was computed as:

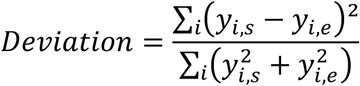

where *i* represents each sample, and *y*_*s*_ and *y*_*e*_ are the simulated and experimental values, respectively. The computed deviation values lie between 0 and 1 and are dimensionless. To compute the overall deviation of simulations from experiments, we summed the deviations from each metric (**Figure S11A**). The simulation with the minimum deviation was considered the best match with experimental data.

Genomic processivity was computed as the average genomic distance that cohesin complexes reached on the simulated polymer, taking into account stalling at CTCF sites (**Figure S10G**). The cohesin loading rate was computed as: *loading rate* = 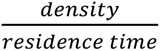 (**Figure S10I**).

##### Comparison of proximal and closed states

We assessed, using simulations, how our segmented proximal state relates to closed states. Closed states were generally identified as proximal states, since 71%, 69%, and 64% of timepoints in the closed states were labelled proximal for L1, L2 and T1, respectively (**Figures S11E** and **S12B**). Proximal states overlapping on closed states generally embraced the closed state and extended before and after closed state starting and ending timepoints, respectively (**Figure S11E**). In addition, proximal states were also identified at non-closed state timepoints (**Figures S11E** and **S12B**). These corresponded to low distances where the two anchors were not directly linked by a single cohesin but the genomic separation between anchors remained low (**Figure S11E**). Because multiple cohesin molecules can simultaneously extrude TADs, anchors can be in spatial proximity even without the presence of a single loop linking them in a closed state (**Figures 5E** and **S11E**). We found that proximal states involved an average of 1.8-2.5 loops, which concomitantly connected TAD anchors (**Figure S12C** and **Video S2**).

##### Statistics

Unless indicated otherwise, we used bootstrap to estimate the standard deviation of our quantifications. To generate bootstrap samples, we randomly selected individual time series with replacements from the full dataset (100% of the dataset size). Normality and homoscedasticity were tested before running any statistical test. The type of statistical test, the sampling size and the type of error bars are indicated in the figure legends.

## Supplementary Figures

**Figure S1:**
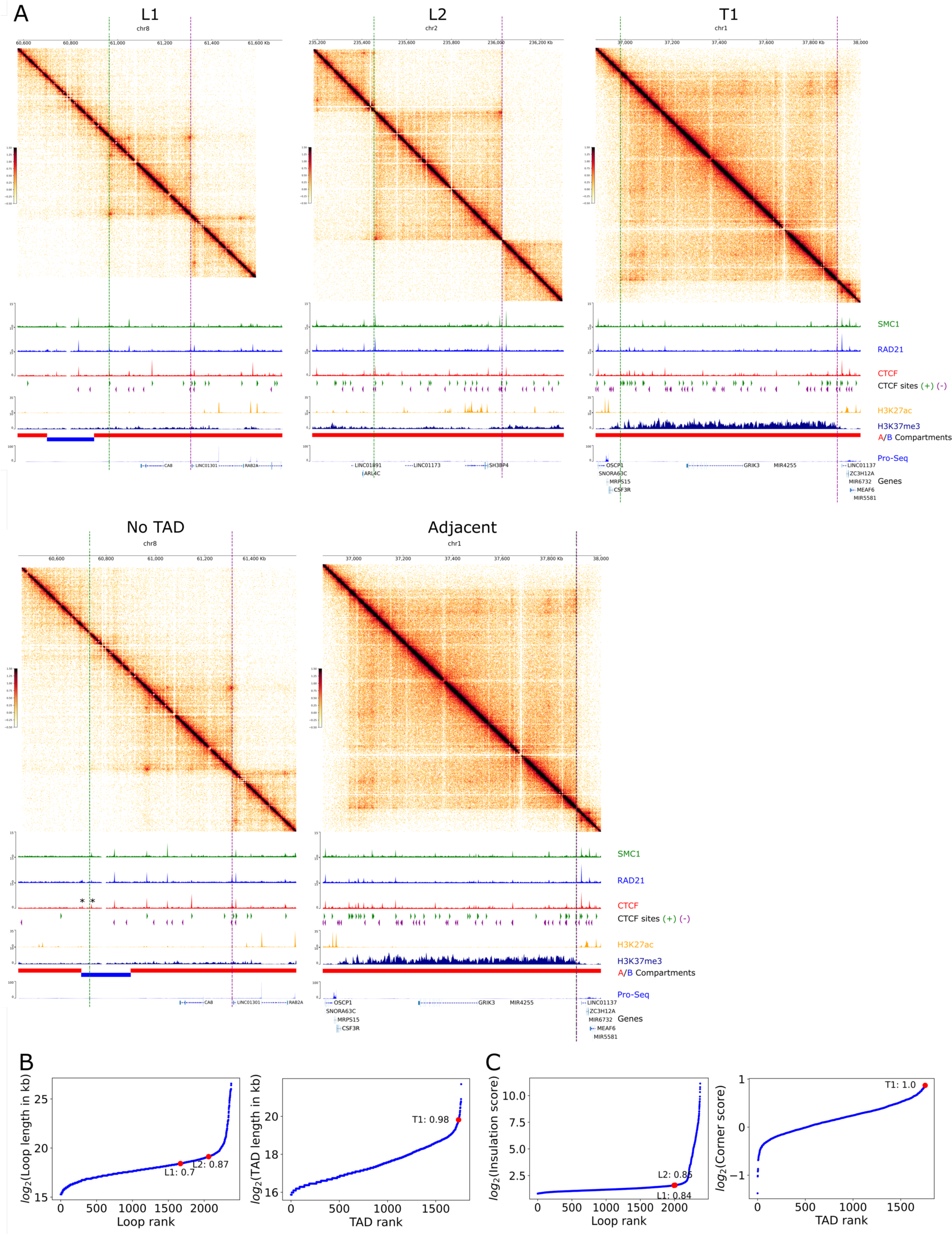
Capture Micro-C maps and genomic features. **A:** Green and magenta dotted lines indicate the genomic location of TetO and CuO repeat array insertion, respectively. Capture Micro-C maps show a 1125 kb region centered on TAD anchors. The Capture Micro-C map generated from the T1 cell line was used to illustrate the Adjacent control. Below contact maps are shown ChIP-Seq and PRO-Seq profiles (data from Rao et al^6^), A/B compartments (computed from Hi-C maps of *Rao et al*^6^), genes and orientation of CTCF sites. * indicates ChIP-Seq peaks overlapping non-significant CTCF sites associated with high q-values (>0.34) in the No TAD control. These two CTCF sites are in divergent orientation as compared to the 3’ TAD anchor and thus not expected to form a loop. **B:** Rank of the genomic lengths of loops (left) and TADs (right). **C:** Rank of insulation score (defined by the ratio of observed to expected bottom left from HICCUPS^106^, left) and corner score (as computed by Arrowhead from juicer^106^, right). In **B-C**, chosen loci are highlighted in red, and their respective percentile is indicated. Only cohesin- and CTCF-dependent domains were considered, *i.e.* loops and TADs exhibiting at least one ChIP-Seq peak of SMC1, RAD21 and CTCF at both domain anchors.

**Figure S2:**
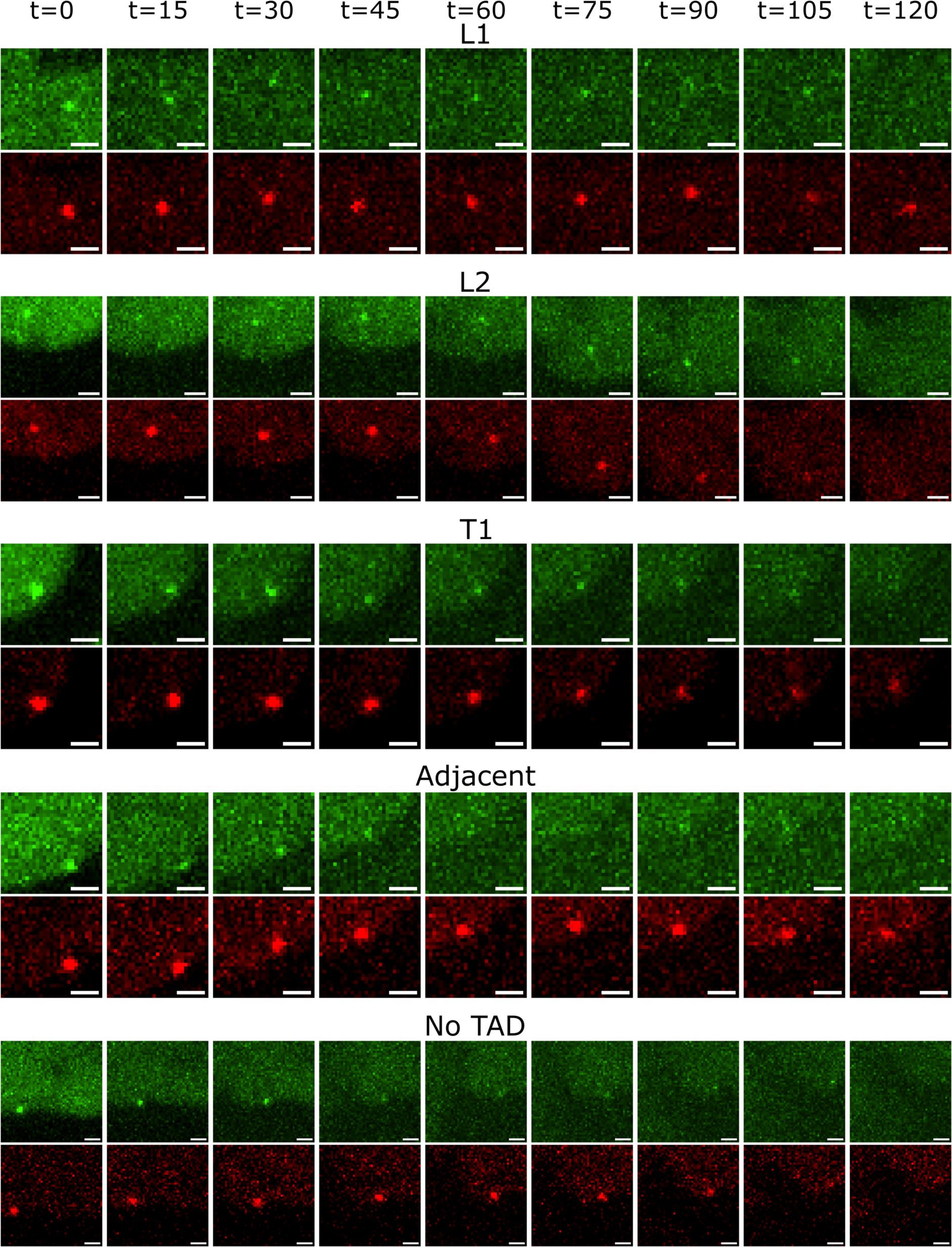
Example time-lapse images. Time-lapse imaging of fluorescently labelled reporters in the two color channels. For visualization, contrast was adjusted independently for each cell line and image intensities were corrected for photobleaching. Maximum intensity projections of 11 z-planes are shown. Timepoints are indicated in minutes. Scale bar: 1 µm.

**Figure S3:**
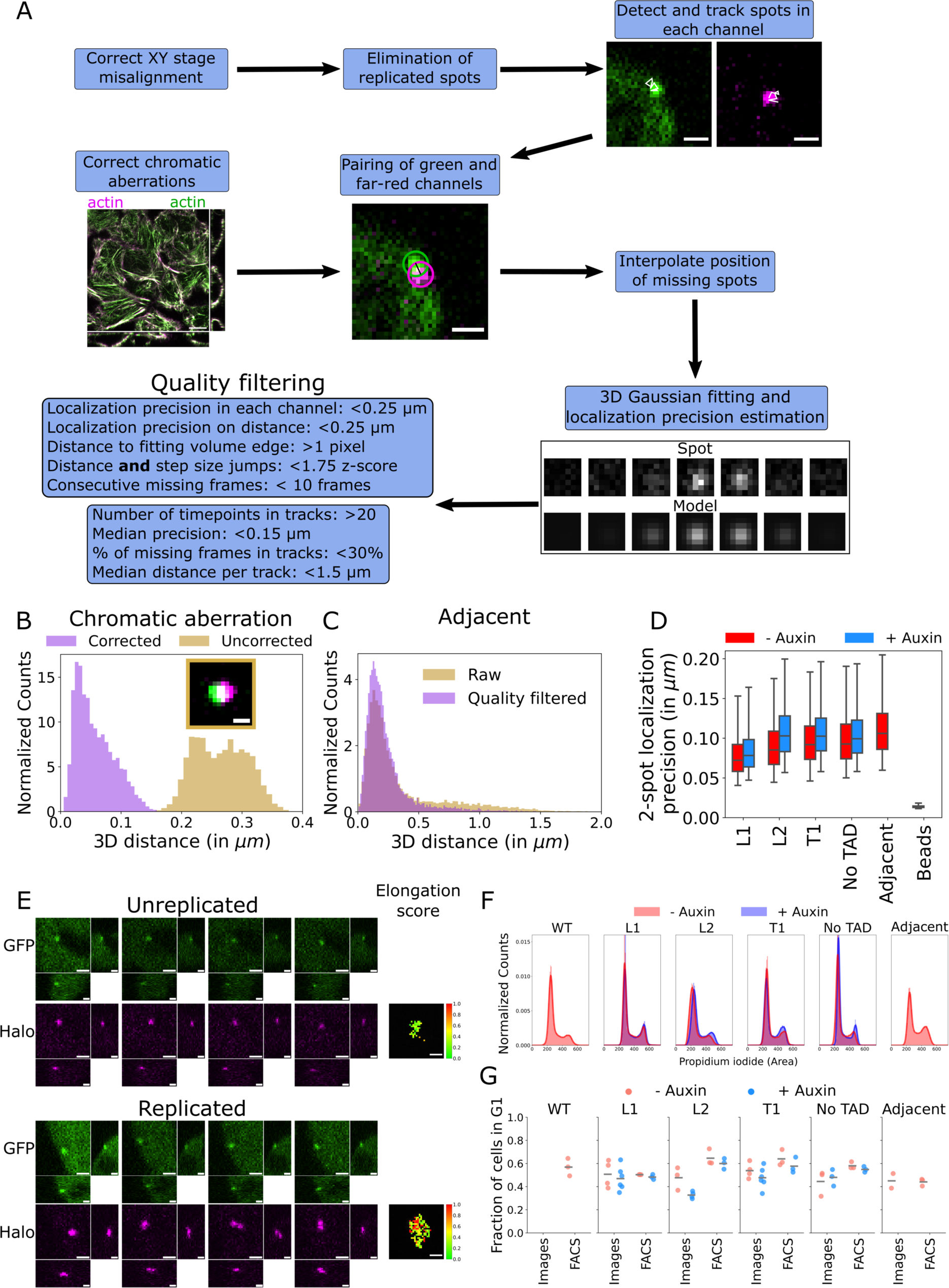
Image analysis pipeline. **A:** Overview of the image analysis pipeline. The actin image is shown without correction of chromatic aberrations. The ‘Spot’ images show different z-planes of the raw image above the corresponding fitted Gaussian model. Scale bar for actin image: 10 µm. Scale bar for fluorescent spot images: 1 µm. **B:** Distance between green and far-red channels of fluorescent beads before and after correction of chromatic aberrations using dual actin labeling to estimate chromatic shifts. The inset shows an image of a fluorescent bead without correction of chromatic aberration. Scale bar: 0.5 µm. **C:** 3D anchor-anchor distance distribution before and after quality filtering in the Adjacent control cell line. **D:** Localization precision of distance measurement estimated from Cramér-Rao bound^50^. Horizontal bars indicate median, lower and upper quartiles. Whiskers extend from 2.5 to 97.5 percentiles. **E-G:** Elimination of cells in S or G2 phases of the cell cycle. **E:** Images of unreplicated (top) and replicated (bottom) fluorescent spots (same contrast for all images) at different timepoints. The spot elongation score was computed to guide replication spot elimination and projected in z and time (right, see Methods). Scale bar: 1 µm. **F:** Distributions of propidium iodide signal assessed by FACS with or without a 3-hour auxin treatment. **G:** Fraction of cells in G1 or early S phase as determined by images or FACS. For images, the fraction of cells remaining after elimination of cells with replicated spots is shown. Each dot shows the median fraction of cells in G1 (or early S phase) across all fields of view acquired during an imaging experiment. N=3 replicates for FACS, N=2-6 replicates for images.

**Figure S4:**
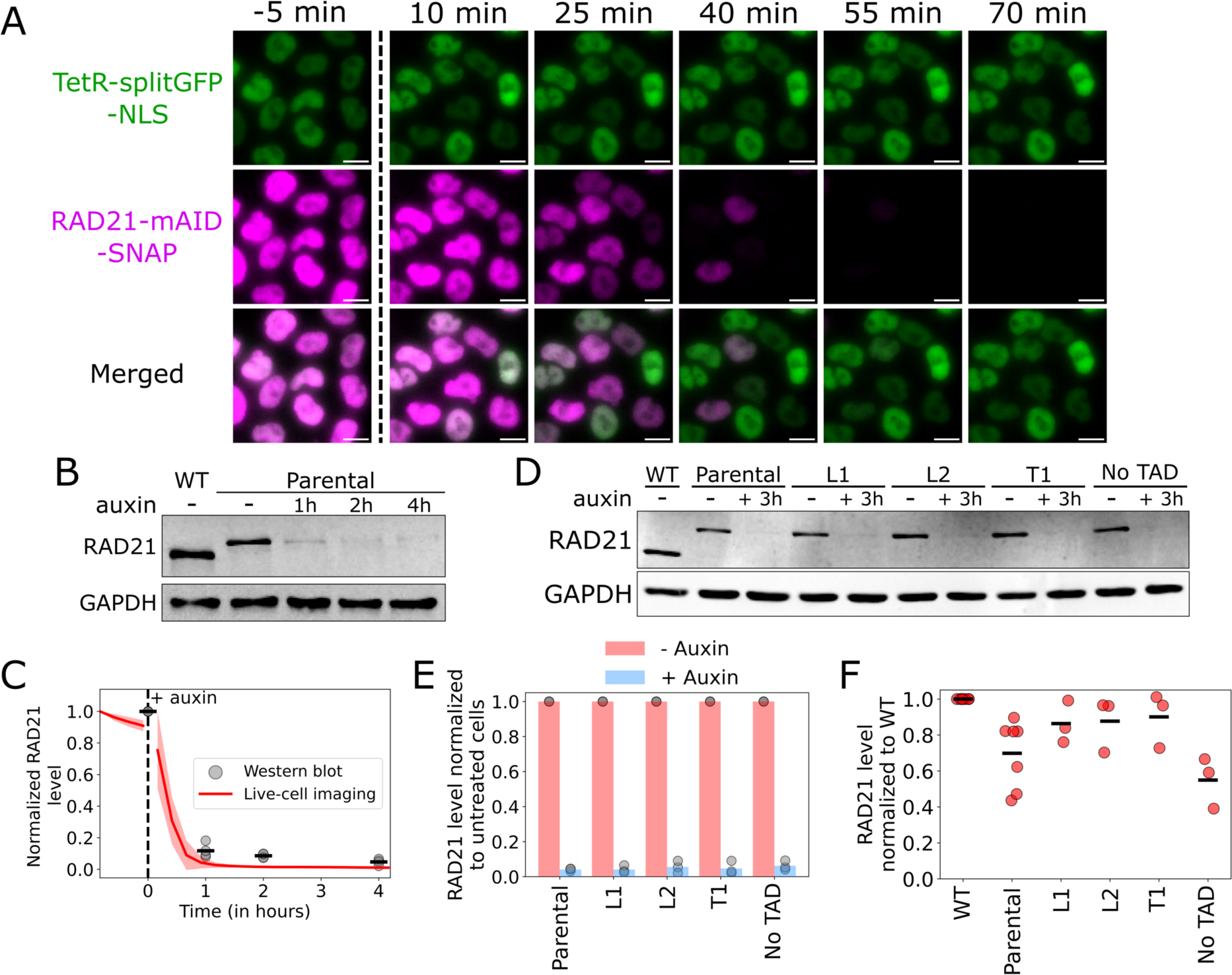
RAD21 auxin-dependent degradation kinetics. **A:** Live-cell images of cell nuclei after auxin treatment. The green signal (TetR-splitGFPx16-NLS) was used to segment nuclei for quantification of RAD21 levels (in **C**). Time after addition of auxin is indicated on top. Scale bar: 10 µm. **B:** Example western blot of RAD21 depletion kinetics upon auxin treatment in the parental cell line. **C:** Quantification of RAD21 level in the parental cell line as function of auxin treatment duration from live-cell imaging (red line) and western blots (grey dots). For live-cell imaging, each timepoint shows the mean normalized RAD21 intensity over all fields of view from N=4 replicates (165-596 cells per replicate). The red shaded area shows the 95% confidence interval. For western blot, horizontal black lines indicate the mean of N=4 replicates. The black vertical dotted line shows the timepoint at which auxin was added to the medium. **D:** Same as **C** for each cell line containing repeat arrays with or without a 3-hour auxin treatment. **E:** Western blot quantification of RAD21 degradation in the different cell lines without (red) or with (blue) a 3-hour auxin treatment. Bars indicate the mean of N=3 replicates, each replicate is shown as a distinct grey dot. **F:** Quantification of basal RAD21 degradation in untreated cell lines as compared to WT RAD21 levels. Black horizontal lines indicate the mean of N=3 replicates (N=7 replicates for WT and parental cell lines).

**Figure S5:**
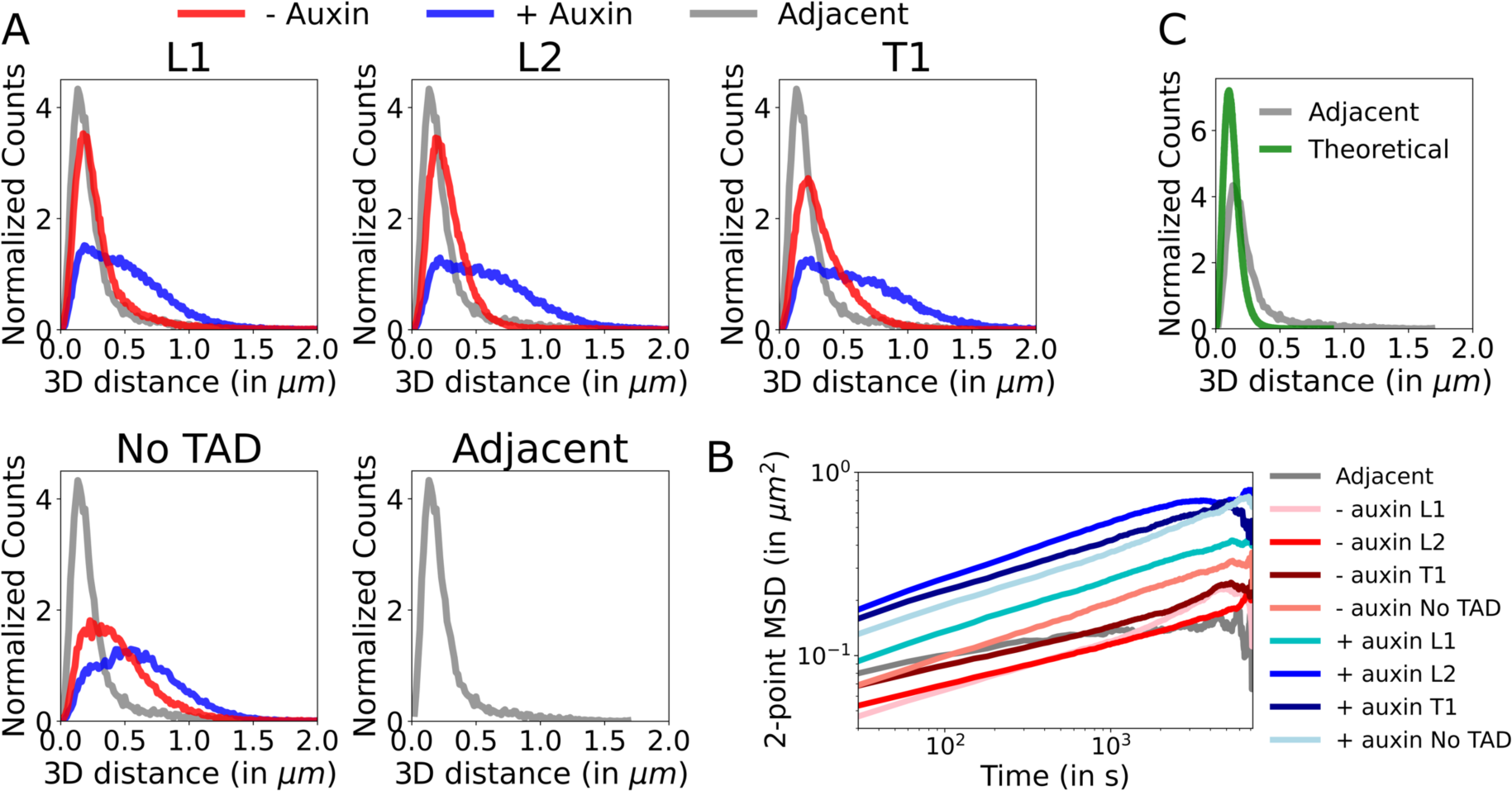
Anchor-anchor distances and motion in untreated or auxin-treated cells. **A:** Histograms of anchor-anchor distances with (blue) or without (red) auxin treatment and the Adjacent control (grey), in which reporters are separated by only 6 kb (grey). **B:** 2-point Mean Squared Displacements (MSD) for untreated (shades of red) and auxin-treated (shades of blue) cell lines, and the Adjacent control (grey). Auxin treatment led to an increase in chromatin mobility. **C:** Anchor-anchor distances of the Adjacent control (grey) and predicted by a theoretical polymer model of closed states (green) assuming 6 kb between reporters, as in the Adjacent control, and taking into account localization errors measured in this cell line.

**Figure S6:**
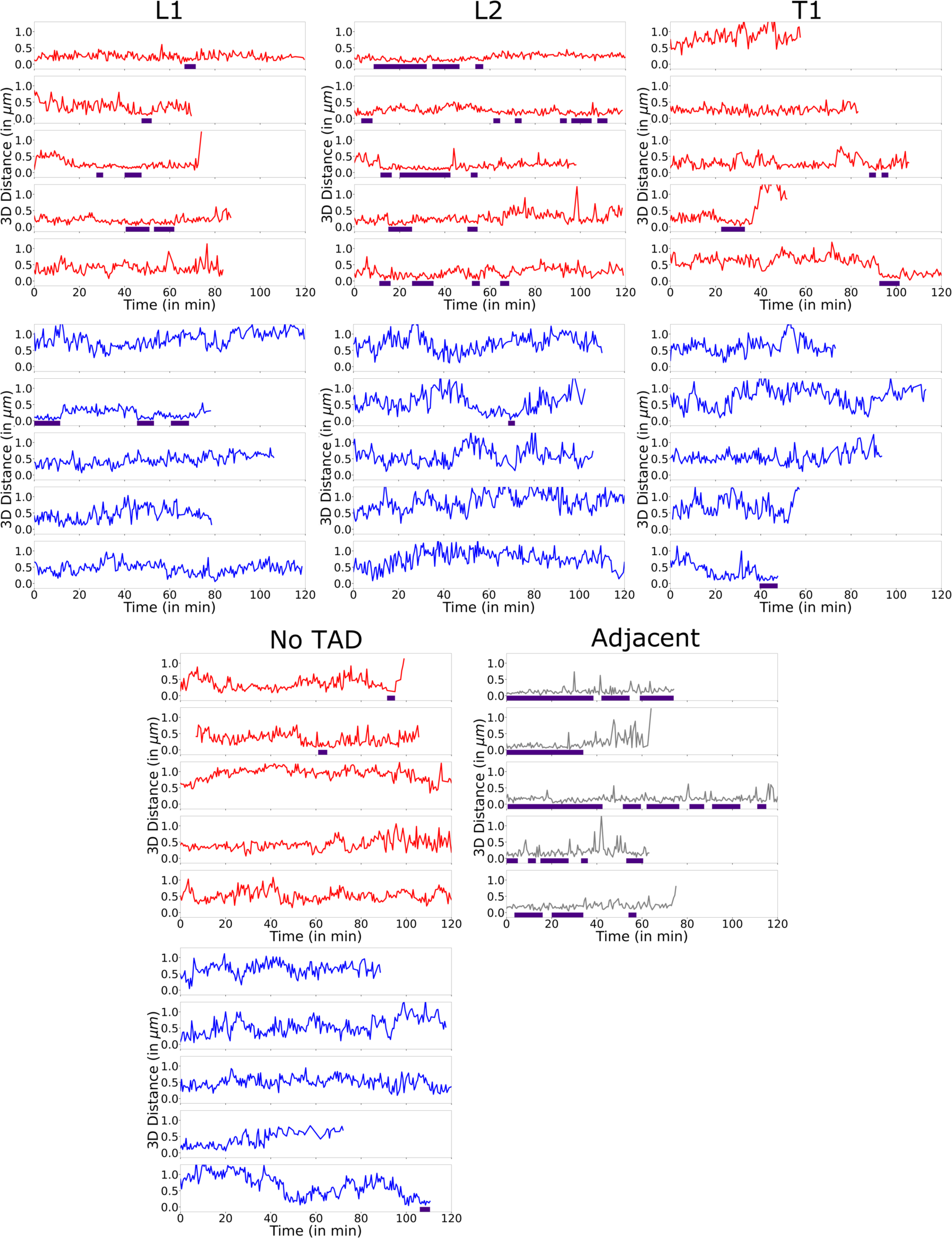
Example time series of anchor-anchor distances and proximal state segmentation. Time series of anchor-anchor distances of untreated (red), auxin-treated (blue) cells and the untreated Adjacent control (grey). Segmented proximal state intervals are indicated by indigo bars.

**Figure S7:**
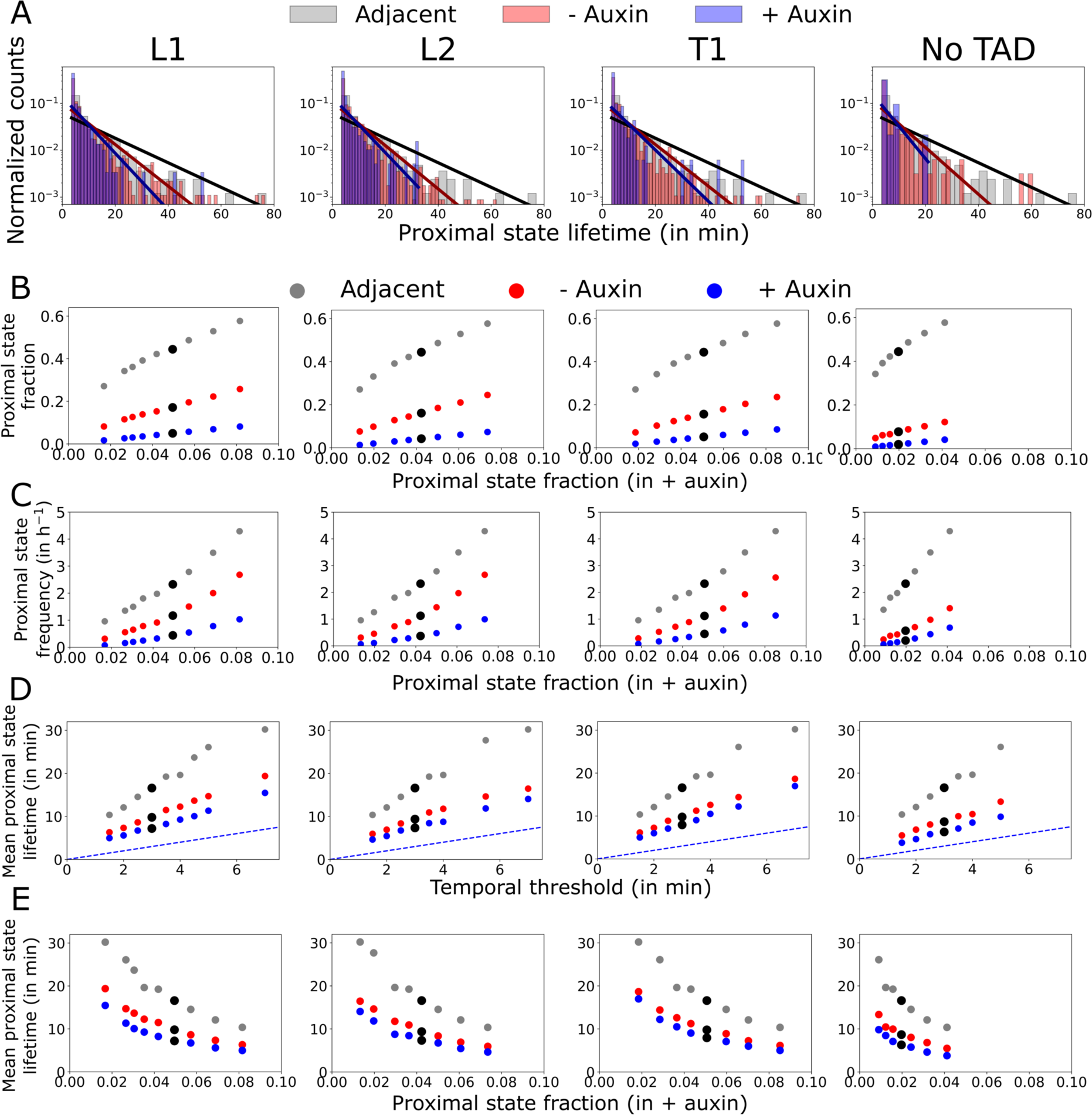
Quantification of proximal states. **A:** Distribution of proximal lifetimes and the corresponding exponential fit (solid lines). **B, C and E:** Fraction of proximal states (**B**), frequency (**C**) and lifetimes (**E**), for various temporal thresholds used in proximal state segmentation, as function of the fraction of proximal states in auxin-treated cells. **D:** Mean proximal state lifetimes estimated as function of temporal thresholds. In **B-E**, each dot corresponds to the estimation resulting from a distinct temporal threshold and the black dot indicates the temporal threshold used to segment proximal states in Figures 2, **3**, **5** and **6**.

**Figure S8:**
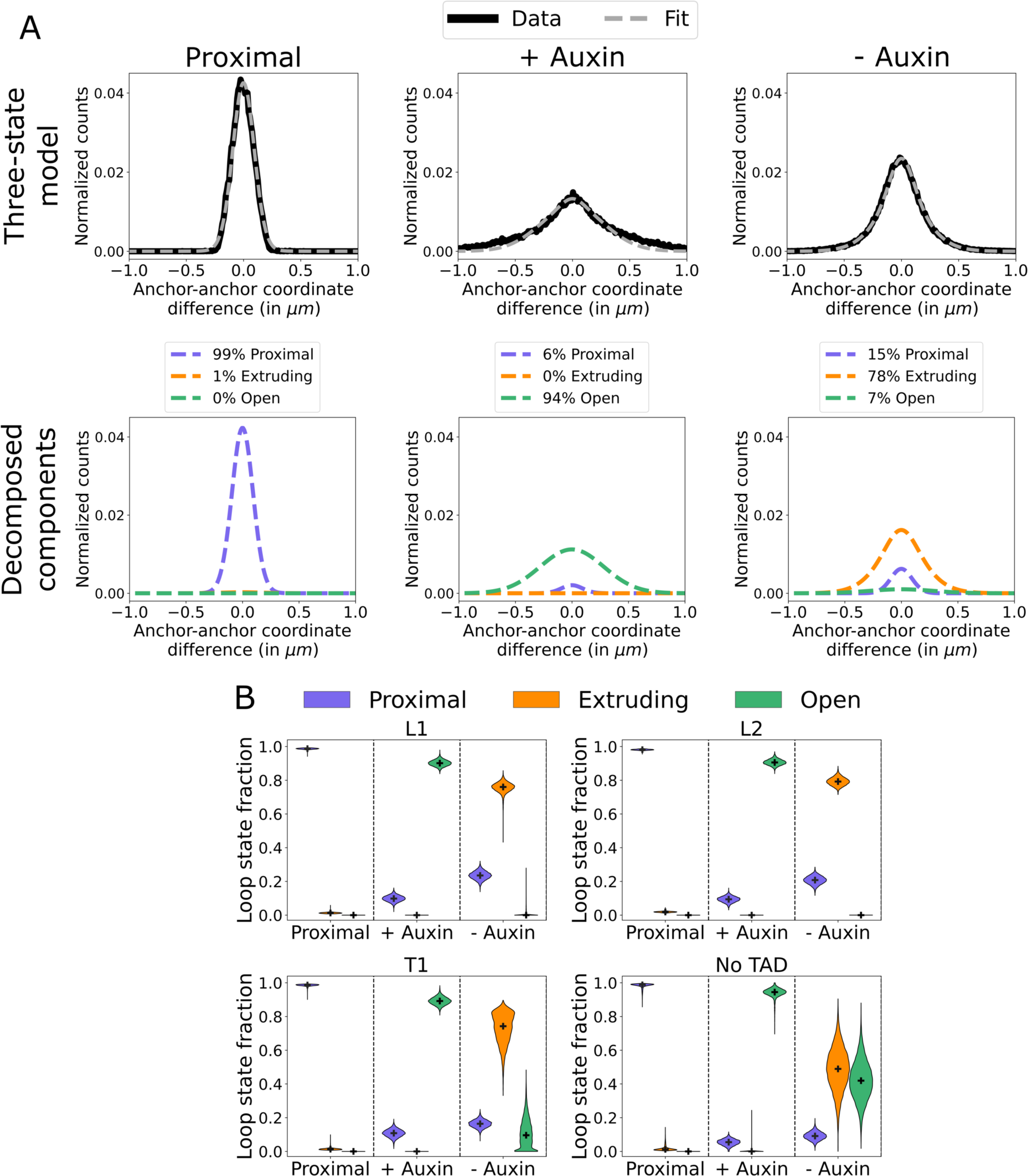
Fractions of loop states estimated from an analytical model of coordinate difference distributions. **A:** Top: Experimental distribution of anchor-anchor coordinate difference (solid line) and the fitted three-state analytical model (dotted line). Bottom: The three components (corresponding to the proximal, extruding and open states) estimated from fitting of the three-state model to the above experimental distribution. The weighted sum of the decomposed components (bottom) yields the three-state model illustrated by the dotted line in the top panel. The T1 TAD was used as an example. **B:** Loop state fractions estimated from the anchor-anchor coordinate difference distributions, in untreated or auxin-treated cells and using only time points segmented as proximal states in untreated cells (‘Proximal’). The black cross indicates the median, violin plots extend from the minimum to maximum. All distributions of loop state fractions are significantly different from each other within each condition (conditions are separated by vertical dashed lines), as assessed by a Kruskal-Wallis test followed by a Dunn’s posthoc test. N=10,000 bootstraps.

**Figure S9:**
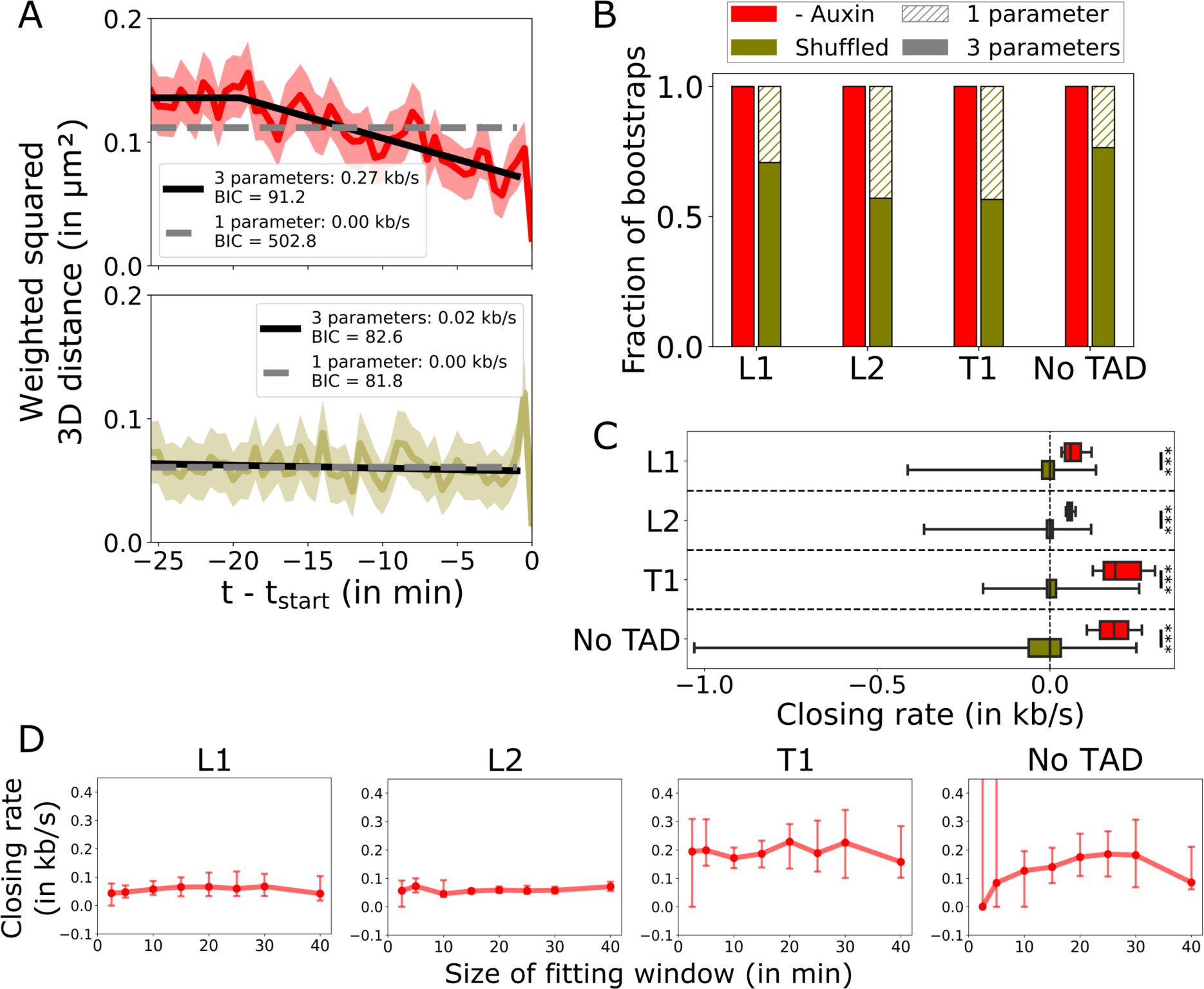
Estimation of closing rate from aligned squared distances. **A:** Squared 3D anchor-anchor distance aligned on *t*_*start*_. The corresponding closing rate fits using a 1-(dashed grey line) or 3-parameter (solid black line) model on untreated (red, N=114) or randomly shuffled (dark yellow, N=79) time series. The BIC value is indicated for each fit. The T1 TAD was used as an example. **B:** Fraction of bootstraps exhibiting the lowest BIC for 1-(hatched) or 3-parameter (filled) fits for untreated (red) or randomly shuffled (dark yellow) time series. **C:** Same as Figure 3E, but with the randomly shuffled time series. The vertical dotted line indicates *x* = 0 kb/s. Boxplot whiskers extend from 10 to 90 percentiles. ***: P-value<0.001 from a Mann-Whitney-U test. In **B** and **C**, N=5,000 bootstraps. **D:** Estimated closing rate as function of the fit window size for untreated cells. Data are represented as mean and error bars extend from the 10 to 90 percentiles of N=5,000 bootstraps.

**Figure S10:**
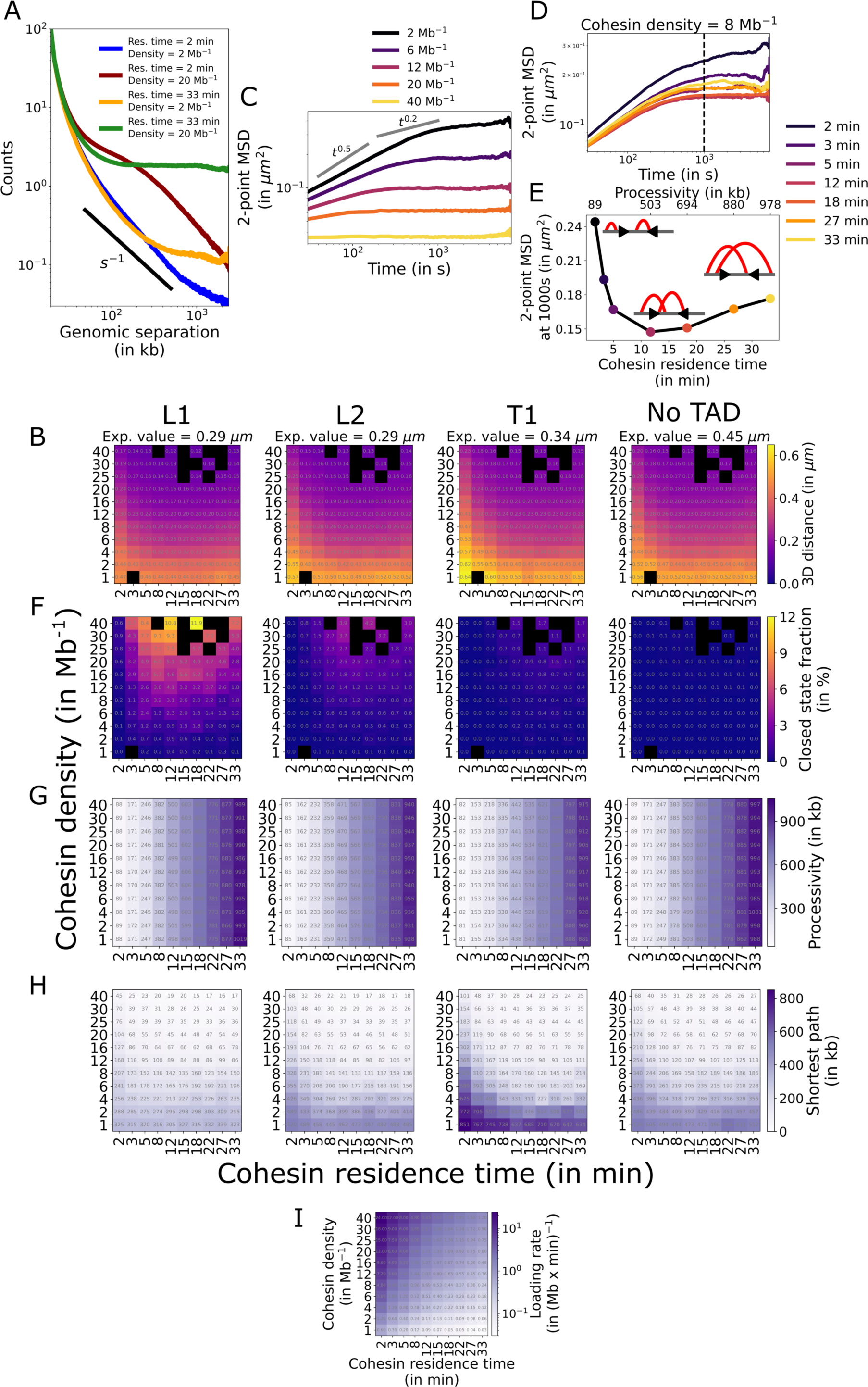
Influence of cohesin residence time and density on TAD dynamics. **A:** Contact counts as function of genomic separation for contact maps of the L1 TAD and cohesin dynamics parameters displayed in Figure 4C. **B and F-I:** Anchor-anchor distance (**B**), closed state fraction (**F**), cohesin processivity (**G**), shortest 1D path connecting TAD anchors (**H**) and cohesin loading rate (**I**) as function of cohesin density and residence time. **C-D:** 2-point MSD curves from polymers with various cohesin densities (**C**) or residence times (**D**), for the L1 TAD. The dotted line in **D** shows the timepoint at which MSD values were observed in **E**. **E:** Absolute 2-point MSD value at 1000 s (dotted line in **D**) depending on cohesin residence time. Schemes represent the length of loops at different cohesin residence times. A cohesin motor speed of 1 kb/s was used in this figure.

**Figure S11:**
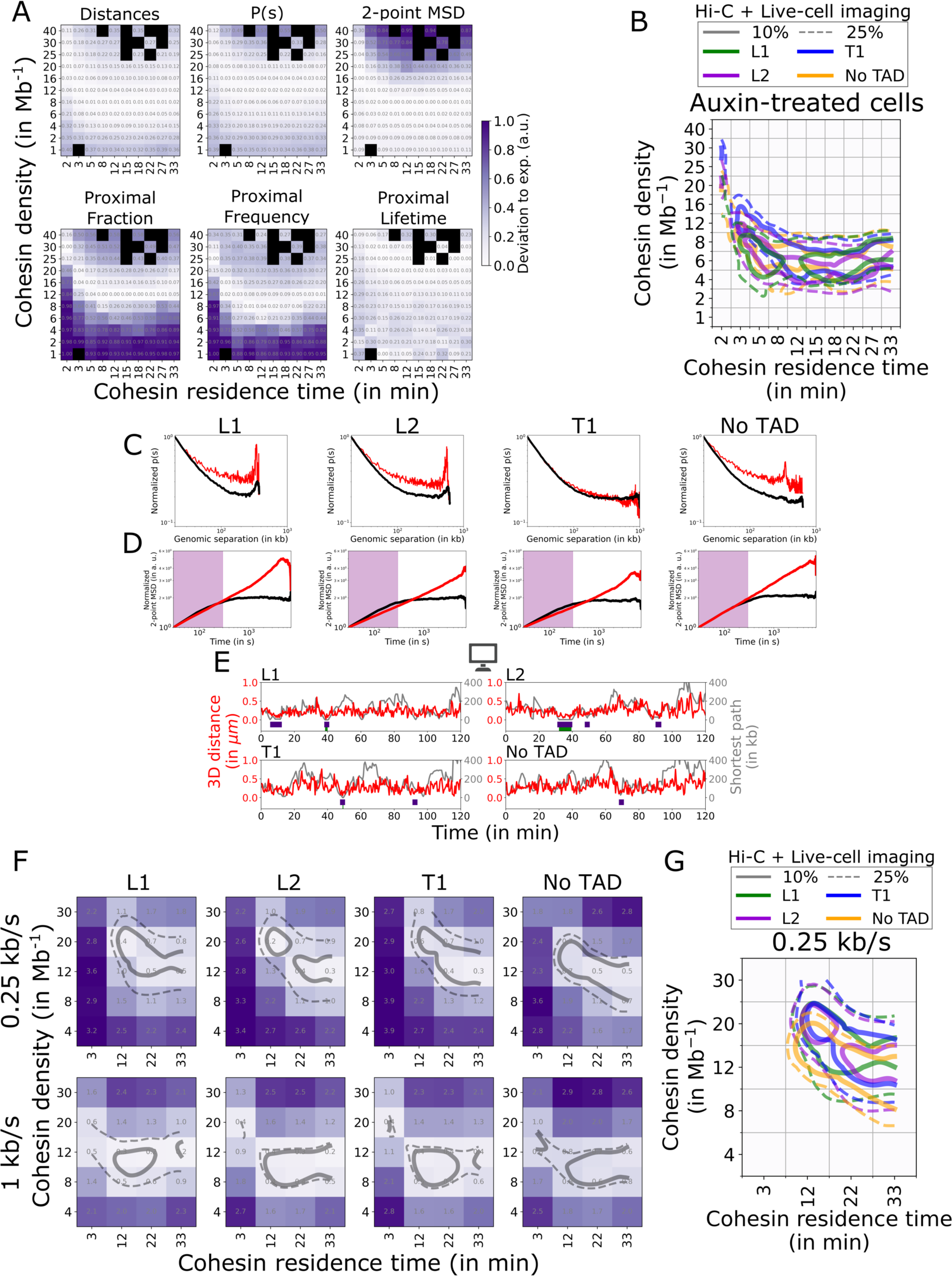
Comparison of simulated and experimental TAD dynamics. **A:** Deviations of polymer simulations from experimentals for each feature of the L1 TAD. The sum of each heatmap yields the deviation map shown in Figure 5A. **B:** Same as Figure 5C, but using experimental data from auxin-treated cells. **C-D:** Simulated (black) and experimental (red) normalized p(s) (**C**) and 2-point MSD (**D**) curves. The simulated curves correspond to the set of simulated parameters represented by a black asterisk in Figure 5C. In **D**, the purple area shows the part of the curve used to compare simulated and experimental curves. **E:** Anchor-anchor distance time series (red) from polymer simulations of the set of parameters represented by a black asterisk in Figure 5C. The shortest 1D path connecting TAD anchors is indicated in grey. Purple and green rectangles at the bottom of time series indicate segmentation of proximal and closed states, respectively. In **A-E**, a cohesin motor speed of 1 kb/s was used. **F:** Deviation of polymer simulations from experiments, as in Figure 5A, but for cohesin motor speeds of 0.25 kb/s (top), results for 1 kb/s are shown for comparison (bottom). **G:** Same as Figure 5C, but for cohesin motor speeds of 0.25 kb/s.

**Figure S12:**
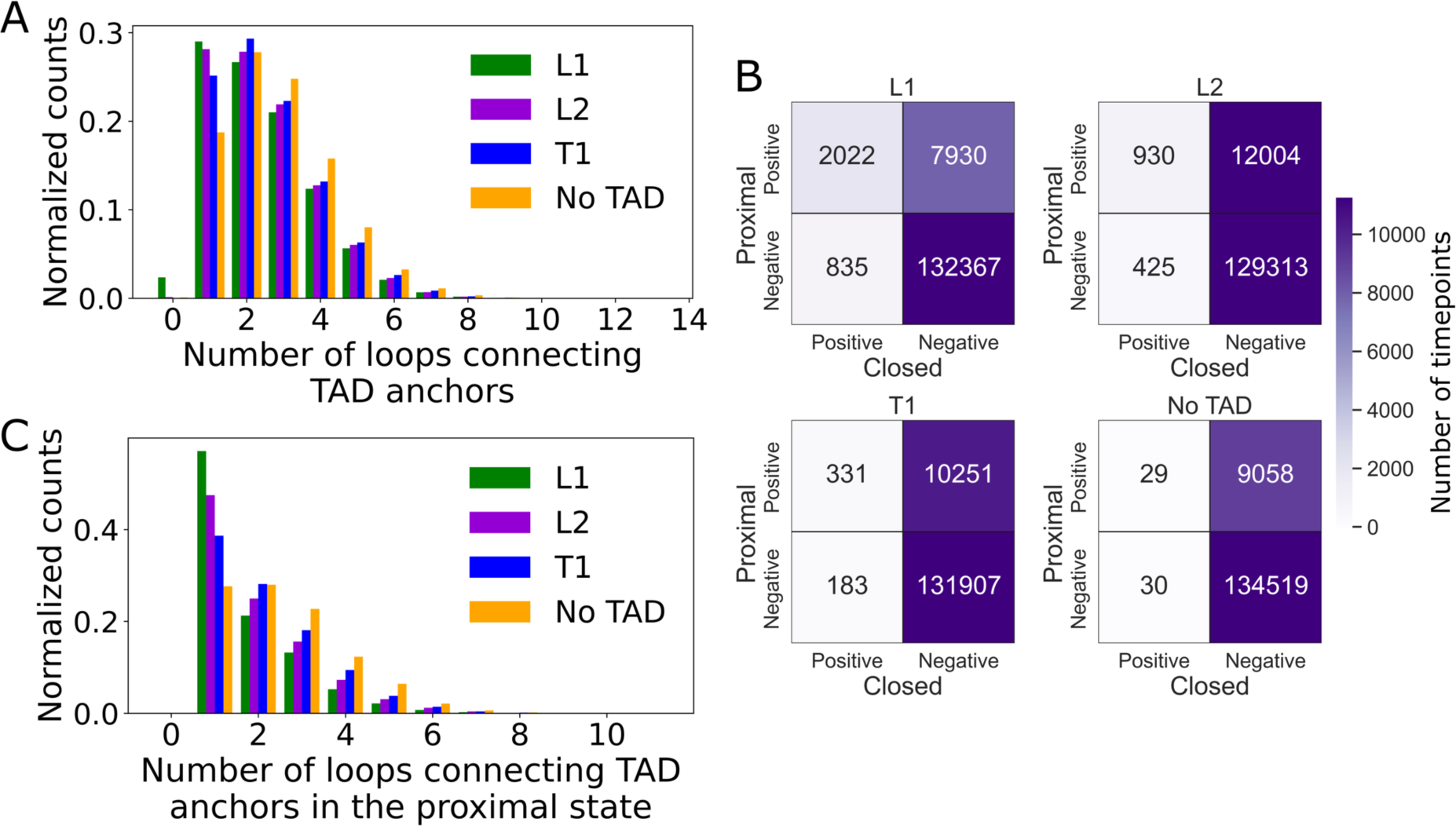
Characterization of the shortest 1D path and proximal states. **A:** Number of loops connecting TAD anchors by the shortest 1D path at any timepoint. **B:** Comparison of closed and proximal states on polymer simulations. The number of timepoints assigned to each class is shown. **C:** Number of loops connecting TAD anchors by the shortest 1D path for proximal states. In **A-C**, the set of simulated parameters highlighted by a black asterisk in Figure 5C and a cohesin motor speed of 1 kb/s were used.

**Figure S13:**
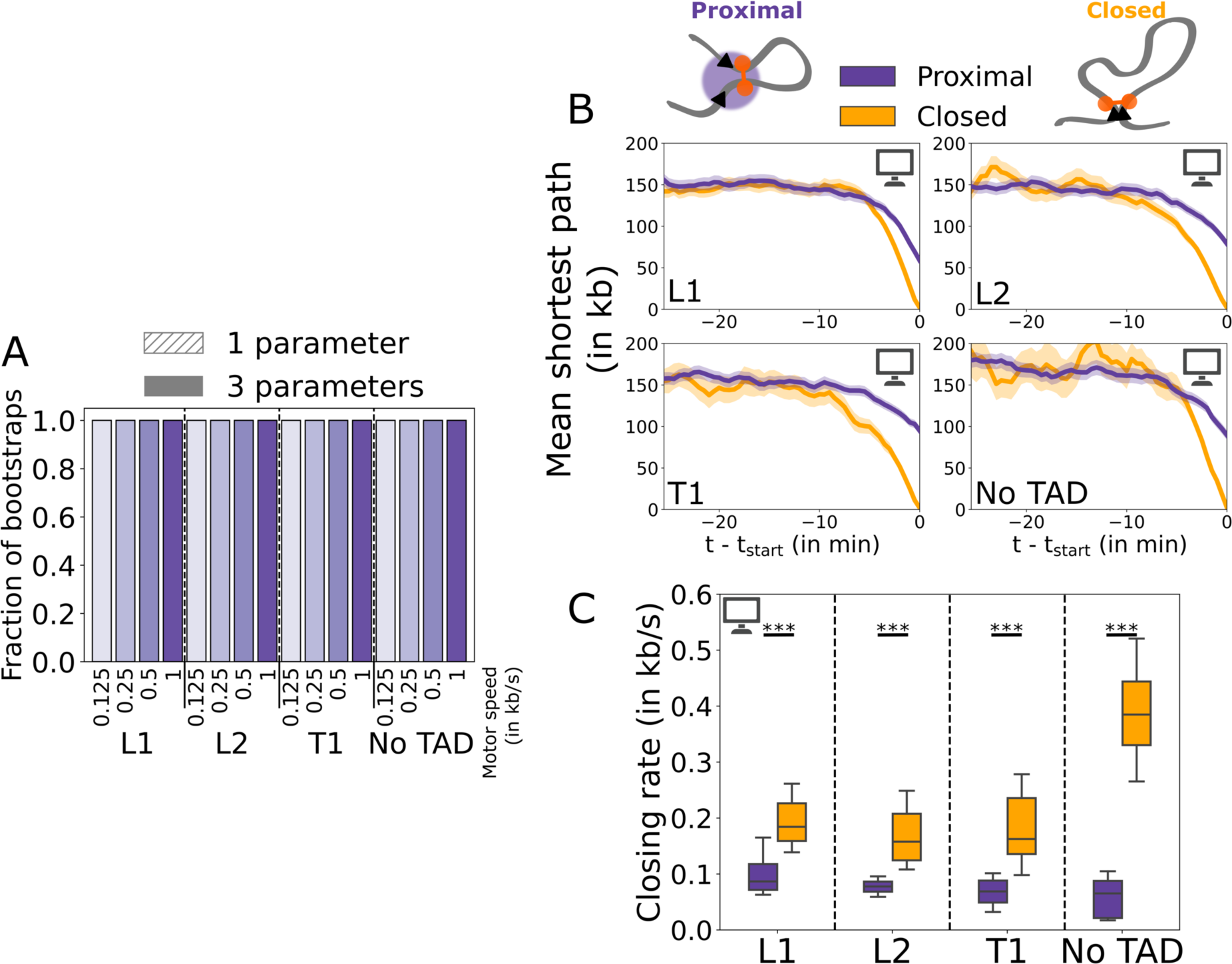
Proximal state segmentation decreases the estimated closing rate. **A:** Fraction of bootstraps consistent with a 1- (filled) or 3-parameter (hatched) model based on the minimal value of BIC, for simulated sets of parameters in Figure 6C. N=5,000 bootstraps. For all sets of parameters, all bootstraps were consistent with a 3-parameter model. **B:** Mean shortest 1D path aligned on proximal (purple) or closed (orange) states. ***: P-value < 0.001 from a Mann-Whitney-U test. **C:** Closing rate estimated from time series aligned on the proximal (purple) or closed (orange) states. A cohesin motor speed of 1 kb/s was used. In **A** and **C**, N=5,000 bootstraps.

**Video S1: Time lapses of the L1 locus.**

Live-cell imaging of L1 TAD anchors from cells left untreated (top) or after a 2-hour auxin-treatment (bottom). Timestamp indicates time as minutes:seconds.

**Video S2: 1D simulations of each genomic region.** A cohesin density of 8 Mb^-1^, residence time of 18 min and motor speed of 1 kb/s were used. Red dotted lines indicate the anchors of each TAD (or fluorescent report for the No TAD control). Timestamp indicates time as hours:minutes.

**Table S1:** Overview of imaging experiments. All statistics are from quality-filtered datasets.

| Condition | Number of movies | Number of time series | Average length of time series (in min) | Median localization error ($\sigma_x, \sigma_y, \sigma_z$) in nm (green channel) | Median localization error ($\sigma_x, \sigma_y, \sigma_z$) in nm (far-red channel) | Median precision of 3D distance (in nm) |
| --- | --- | --- | --- | --- | --- | --- |
| L1 -auxin | 30 | 538 | 73 | 24, 25, 46 | 16, 16, 32 | 72 |
| L1 +auxin | 20 | 470 | 75 | 25, 27, 49 | 18, 18, 37 | 78 |
| L2 -auxin | 50 | 531 | 79 | 29, 33, 59 | 15, 15, 31 | 85 |
| L2 +auxin | 32 | 427 | 72 | 36, 41, 74 | 16, 16, 34 | 103 |
| T1 -auxin | 49 | 694 | 61 | 33, 37, 69 | 12, 13, 27 | 92 |
| T1 +auxin | 40 | 393 | 49 | 36, 40, 75 | 15, 16, 33 | 103 |
| No TAD -auxin | 27 | 252 | 77 | 30, 34, 61 | 19, 20, 40 | 93 |
| No TAD +auxin | 14 | 167 | 78 | 31, 35, 63 | 22, 22, 45 | 99 |
| Adjacent -auxin | 25 | 150 | 47 | 38, 44, 80 | 14, 14, 29 | 106 |

**Table S2:** Distances between repeat array center and CTCF sites defining the loop anchor in kb.

|  | L1 | L2 | T1 | No TAD |
| --- | --- | --- | --- | --- |
| Distance from repeat array center to 5' anchor (in kb) | 2.3 | 2.3 | 3.3 | 0 |
| Distance from repeat array center to 3' anchor (in kb) | 3.9 | 3.8 | 4.7 | 3.9 |
| Spatial threshold (in $\mu\text{m}$) | 0.199 | 0.218 | 0.236 | 0.220 |

## Notes

### Competing Interest Statement

The authors have declared no competing interest.

